# Data-driven inference of local behavioural rules predicts emergent properties of *Phytophthora* zoospore dispersal

**DOI:** 10.64898/2026.08.12.744352

**Authors:** Joëlle Le Berre, Agnès Attard, Edouard Evangelisti

**Affiliations:** Institut Sophia Agrobiotech, UMR INRAE 1355, Université Côte d’Azur, Sophia Antipolis, France

**Keywords:** oomycete, zoospore, agent-based modelling, hidden Markov model, prediction, microbial dispersal

## Abstract

Relating local behavioural rules to emergent population-level dynamics is a central challenge in quantitative biology. Data-driven agent-based models offer a powerful route to infer such rules directly from experimental observations, but their predictive value ultimately depends on whether the inferred behaviour remains valid beyond the conditions used for model construction. Here, we address this question using biflagellate zoospores of plant-pathogenic *Phytophthora* as an experimentally tractable model system of microswimmers. From nearly 60,000 freely swimming *P. nicotianae* trajectories, we inferred local behavioural rules governing speed and turning, including the temporal persistence of distinct movement states, and implemented them in generative agent-based simulations. An empirical two-state model reproduced the principal emergent properties of experimental dispersal, while independent hidden Markov modelling recovered a closely related behavioural organisation. We then challenged these inferred swimming rules in a distinct biological and physical context, using *P. palmivora* zoospores navigating an obstacle-filled environment. Without recalibrating the underlying swimming behaviour, the model reproduced emergent dispersal when combined with simple collision rules and identified transient post-collision slowdown as a key response required to account for the observed trajectories. These results show that local behavioural rules inferred from individual motion can retain predictive power beyond the conditions from which they were learned. More broadly, our framework provides a way to distinguish models that merely reproduce their training observations from those that capture transferable features of biological movement.

## Introduction

Complex biological spatial dynamics can emerge from local interactions among individual cells or organisms, resulting in collective behaviour at a larger scale (Ouellette & Gordon, 2021). Relating local behaviours to population-level properties is therefore a prominent challenge in quantitative biology (Binny *et al*., 2015). Agent-based models (ABMs) constitute an elegant framework for addressing this problem. In contrast to approaches that directly assign collective properties to a population, ABMs explicitly simulate individual entities’ behaviours and their interactions. The collective properties of the population emerge from the repeated application of local rules governing individual behaviour (Bonabeau, 2002). Such approaches have been successfully used to investigate microbial communities, cell migration, biofilm development, and other systems in which local interactions generate spatially and temporally organised dynamics (Buttenschön & Edelstein-Keshet, 2020; Nagarajan *et al*., 2022).

A central question in ABM approaches is where the rules assigned to individual agents originate. In many mechanistic models, behavioural rules derive from biological knowledge or physical assumptions, while unknown parameters are subsequently calibrated against experimental observations (Sivakumar *et al*., 2022; Cogno *et al*., 2024). Experimental data can also be used to constrain, calibrate or directly infer agent behaviour. For example, data-driven agent-based modelling of the bacterium *Myxococcus xanthus* used an iterative process combining single-cell tracking with simulations to identify behavioural features necessary and sufficient to reproduce the observed aggregation dynamics (Cotter *et al*., 2017). More recently, machine-learning methods have been applied to cell trajectories to infer interaction rules underlying collective cell migration (LaChance *et al*., 2022), and multi-agent neural-network approaches have been developed to learn cellular interaction dynamics from observations (Rocha *et al*., 2025). Systematic workflows have also been developed to calibrate microbial ABMs against time-lapse microscopy data (Ahmadi *et al*., 2024). These advances illustrate a broader shift from predominantly prescribed local rules towards models in which agent behaviour is increasingly constrained or learned from observations.

While inferring rules that reproduce an observed system is an important step in quantitative biology, a stronger test is whether these rules retain predictive power beyond the conditions from which they were inferred (Windrum *et al*., 2007). A model calibrated on a given dataset can accurately reproduce the statistics or emergent properties of that dataset while remaining dependent on the particular conditions under which its parameters were obtained (Gallagher *et al*., 2021). The critical question is therefore whether behavioural rules inferred in one context retain explanatory and predictive power when the physical or biological environment is changed. This distinction is especially important for data-driven models, for which agreement with the training observations may demonstrate successful inference without necessarily establishing that the inferred rules capture transferable features of the underlying behaviour (Kaul & Ventikos, 2015; Yates *et al*., 2018). Determining how far local rules can be transferred beyond the conditions from which they were learned is therefore central to developing genuinely predictive models of biological movement.

Motile microorganisms offer an experimentally tractable system to address this question. Their trajectories can be recorded at single-cell resolution while environmental conditions can be independently manipulated, thereby enabling local behavioural dynamics to be linked to emergent dispersal (Son *et al*., 2015). Among these microorganisms, biflagellate zoospores produced by plant-pathogenic oomycetes of the genus *Phytophthora* provide a particularly favourable experimental system. Species of this genus include major crop pathogens responsible for destructive diseases worldwide, making zoospore dispersal relevant not only as a tractable model of microbial movement but also as a critical step in plant infection (Wang *et al*., 2026). Zoospore motility is essential for dispersal and host targeting and responds to chemical, electrical and physical environmental cues (Judelson & Blanco, 2005; Bassani *et al*., 2020; Kasteel *et al*., 2023). At the same time, large numbers of freely swimming zoospores can be imaged over short experimental timescales, enabling the acquisition of sufficiently large trajectory datasets to infer behavioural statistics at high temporal resolution (Galiana *et al*., 2019; Tran *et al*., 2022; Lupatelli *et al*., 2023; Kasteel *et al*., 2024; Cohen *et al*., 2025). Their swimming environment can also be experimentally altered, providing an opportunity to challenge inferred behavioural rules under conditions distinct from those used for their estimation.

Here, we use *Phytophthora* zoospores to ask whether local behavioural rules inferred directly from individual trajectories can predict emergent dispersal beyond the conditions in which they were learned. From nearly 60,000 freely swimming *P. nicotianae* (syn. *P. parasitica*) trajectories, we first quantify local swimming dynamics and use them to construct generative agent-based models in which behavioural rules are inferred directly from experimental observations. We independently examine behavioural organisation using empirical state classification and hidden Markov modelling, and assess whether these local descriptions reproduce population-level dispersal properties that were not explicitly prescribed to the model. Finally, we challenge the inferred swimming rules in an independent, obstacle-filled environment using *P. palmivora*, without recalibrating the underlying swimming behaviour. This framework therefore allows us to distinguish reproduction of the data used for inference from predictive transfer to a distinct biological system and physical environment.

## Materials and Methods

### Microbial strains and cultivation conditions

*P. nicotianae* INRA-310 was originally isolated from tobacco in Australia. *P. palmivora* P16830 (LILI) was isolated from oil palm in Colombia. Both species are maintained in the *Phytophthora* collection at INRAE, Sophia Antipolis (France). Mycelium was cultivated on V8 agar at 24°C in the dark (*P. nicotianae*) or 25°C under constant light (*P. palmivora*). For *P. nicotianae*, sporulation was induced by transferring fragmented mycelium from one-week-old plates onto water agar plates, followed by a four-day incubation at 24°C under a 16 h photoperiod. *P. palmivora* sporangia formed spontaneously after one week of cultivation on V8 agar. Zoospores of both species were released by incubating sporulating cultures at 4°C for 30 min, followed by flooding with sterile water. Zoospore concentration was determined using a hemocytometer and adjusted to the desired concentration for subsequent experiments.

### Image acquisition

Zoospore suspensions were diluted to 10⁶ zoospores per millilitre in sterile water. A 10 µL droplet was introduced into a water-filled observation chamber and immediately imaged using an Axio Zoom microscope (Zeiss, Germany) equipped with a PlanNeoFluar Z 1.0× objective (zoom factor 1.2×) operated in dark-field illumination mode, at a final magnification of 12.5×. Images were acquired at 13.3 frames per second for 1 min using a Hamamatsu ORCA-Flash4.0 LT camera. This frame rate maintained a mean inter-frame displacement of less than 20 µm, thereby facilitating trajectory reconstruction. The resulting time-lapse series, consisting of approximately 800 frames (1 minute), were used directly for image preprocessing and subsequent trajectory analysis.

For imaging in crowded environments, zoospores were allowed to swim in a 0.17-mm-deep water-filled chamber. Ten-micrometre Polybead polystyrene microspheres (Polysciences, USA) were then added and allowed to sediment onto the bottom surface, forming a stable obstacle field. A droplet containing zoospores was subsequently added at the edge of the chamber, allowing cells to progressively enter the bead-containing region. Zoospores were allowed to explore the crowded environment for 5 min before image acquisition.

### Image pre-processing

Minimal image pre-processing was performed before trajectory reconstruction. Time-lapse frames were imported as an image sequence. Out-of-focus halos were first corrected across the stack using the ImageJ Subtract Background algorithm with a rolling-ball radius of 10 pixels and the sliding-paraboloid option enabled. A temporal median projection was then calculated across the complete stack to estimate the persistent background component. This median image was subtracted from every frame to reduce stationary structures and residual background signal. Negative pixel values resulting from subtraction were clipped to zero. Image intensities were normalised using a fixed window (179 intensity units) centred at a level of 77. This same intensity transformation was applied uniformly to all frames before export as a multi-frame TIFF file for subsequent trajectory reconstruction in TrackMate.

### Trajectory reconstruction with TrackMate

Time-lapse microscopy stacks were analysed with TrackMate 7 (<u>imagej.net/plugins/trackmate/</u>) (Ershov *et al*., 2022). Zoospores were detected using the Laplacian of Gaussian (LoG) detector, with an estimated particle diameter of 10 µm, a quality threshold of 80, and sub-pixel localisation enabled. Median filtering within the TrackMate detector was disabled. Trajectories were reconstructed using the Sparse Linear Assignment Problem (LAP) tracker with a maximum linking distance of 100 µm. Gap closing was enabled over a maximum distance of 10 µm and a maximum temporal gap of one frame. No additional spot or track filters were applied after reconstruction, and all available TrackMate feature analysers were executed. For crowded-environment experiments only, trajectories with a net displacement below 20 µm were excluded to minimise residual detections originating from stationary beads that were not completely eliminated during image pre-processing. Spatial and temporal calibration values were read directly from the image metadata, allowing positions, distances and speeds to be expressed in physical units. For each image sequence, the complete TrackMate model and analysis settings were saved as an XML file. For quality control and visual inspection, trajectories were displayed on a black background. At each frame, trajectory segments ending during the preceding 30 frames were drawn as cyan lines with progressively decreasing intensity. Detected zoospores in the current frame were represented as filled magenta discs. This representation facilitated rapid inspection of trajectory continuity, tracking artefacts, and overall tracking accuracy.

### Automatic trajectory analysis

All trajectory analyses were performed using a fully automated Python workflow developed for this study. The complete workflow is distributed together with shell scripts that automatically create the required Python virtual environment, install all dependencies, and execute the analysis pipeline. The workflow was developed and tested with Python 3.13 (www.python.org) (van Rossum, 2025). Starting from the XML files exported by TrackMate, the pipeline reconstructed individual trajectories, computed global trajectory descriptors and local behavioural variables, generated graphical outputs, estimated the parameters required for empirical agent-based simulations, and performed hidden Markov modelling. Global descriptors characterised the geometry and exploratory behaviour of complete trajectories, whereas local descriptors quantified instantaneous swimming behaviour at the level of individual displacements. Local measurements were used for behavioural inference and model calibration, while global descriptors provided independent benchmarks for model evaluation.

#### Global trajectory descriptors

Trajectory-level descriptors, including straightness, net displacement, and mean squared displacement (MSD), were calculated from the reconstructed trajectories **(Table 1)**. These descriptors characterise emergent properties of zoospore movement and were not used during model calibration, thereby providing independent benchmarks for model evaluation.

**Table 1.** Global trajectory descriptors used for model evaluation. Global descriptors were calculated from complete reconstructed trajectories and were excluded from model calibration.

| Descriptor | Definition | Unit | Interpretation |
| --- | --- | --- | --- |
| Straightness | $\frac{\ \mathbf{x}_N - \mathbf{x}_1\ }{\sum_{i=1}^{N-1} \ \mathbf{x}_{i+1} - \mathbf{x}_i\ }$ | — | 1: perfectly straight trajectory; 0: highly tortuous trajectory |
| Net displacement | $D = \ \mathbf{x}_N - \mathbf{x}_1\ $ | $\mu\text{m}$ | Straight-line distance between first and last positions. |
| Mean squared displacement (MSD) | $\text{MSD}(\tau) = \left\langle \ \mathbf{x}(t + \tau) - \mathbf{x}(t)\ ^2 \right\rangle_t$ | $\mu\text{m}^2$ | Average squared displacement as a function of time lag. |

#### Extraction of local behavioural descriptors

Local behavioural descriptors, including swimming speed, turning angle, speed autocorrelation, turning autocorrelation, and speed-turn coupling, were quantified from the reconstructed trajectories **(Table 2)**. These measurements were subsequently used to infer the empirical distributions and statistical dependencies governing local swimming behaviour. Together, they provided the experimental basis for behavioural-rule inference and agent-based modelling.

**Table 2.**
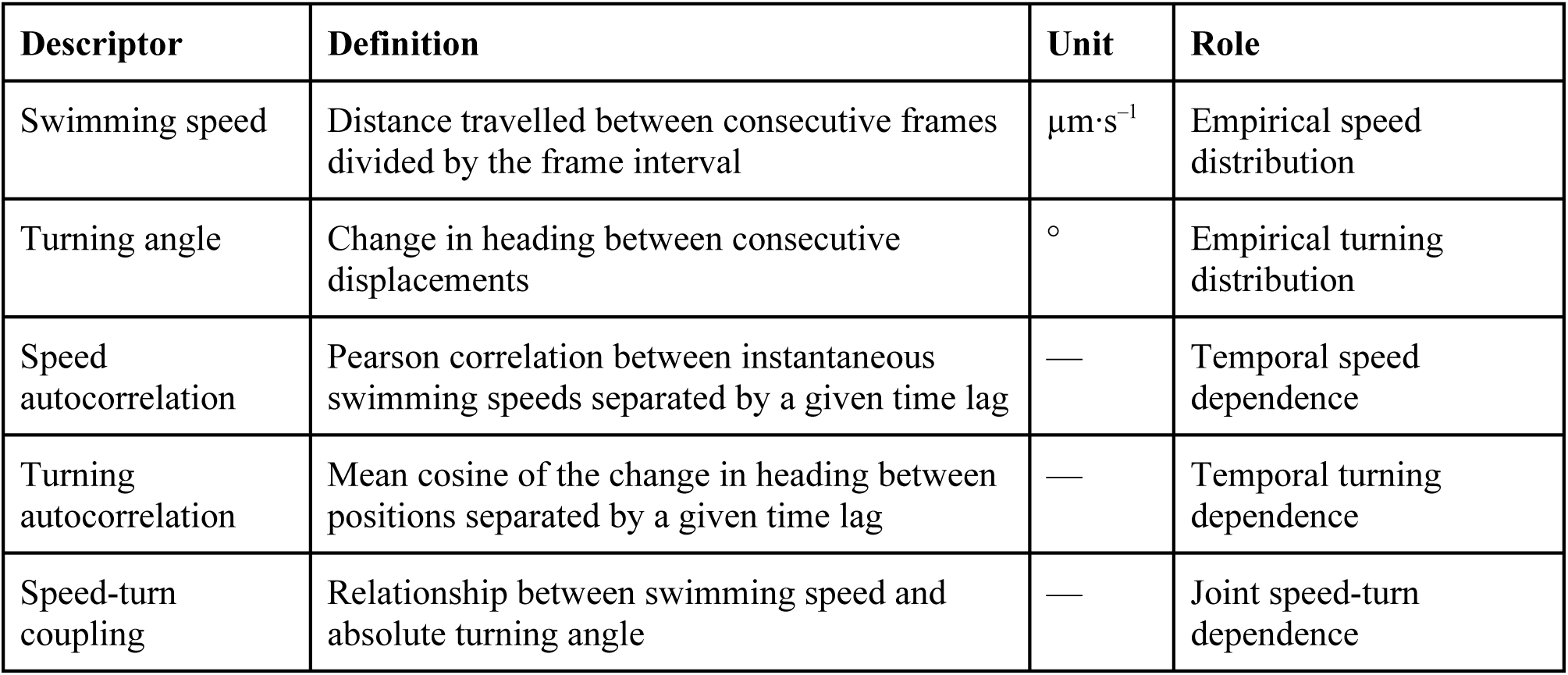
Local behavioural descriptors and derived behavioural parameters. Local behavioural descriptors were calculated from reconstructed trajectories. These measurements and their derived statistics were used to infer the local behavioural rules implemented in the agent-based cellular automata (ABCA) model.

| Descriptor | Definition | Unit | Role |
| --- | --- | --- | --- |
| Swimming speed | Distance travelled between consecutive frames divided by the frame interval | $\mu\text{m}\cdot\text{s}^{-1}$ | Empirical speed distribution |
| Turning angle | Change in heading between consecutive displacements | $^{\circ}$ | Empirical turning distribution |
| Speed autocorrelation | Pearson correlation between instantaneous swimming speeds separated by a given time lag | — | Temporal speed dependence |
| Turning autocorrelation | Mean cosine of the change in heading between positions separated by a given time lag | — | Temporal turning dependence |
| Speed-turn coupling | Relationship between swimming speed and absolute turning angle | — | Joint speed-turn dependence |

#### Hidden Markov modelling

To investigate whether local zoospore dynamics could be parsimoniously described by a limited number of hidden behavioural states, Gaussian hidden Markov models (HMMs) with diagonal covariance matrices were fitted using instantaneous swimming speed, absolute turning angle, and signed acceleration as emission variables. To reduce the influence of long-tailed distributions, speed and absolute turning angle were log-transformed as log(1 + *x*), while signed acceleration was transformed as asinh(*x*/100). The three transformed variables were subsequently standardised before model fitting. Individual trajectories were treated as independent sequences, and only trajectories containing at least 10 usable observations were included. Models were fitted for a maximum of 500 iterations, with a convergence tolerance of 10⁻⁴ and a minimum covariance of 10⁻⁴. Models containing two to seven hidden states were fitted using ten independent initialisations for each number of states, with the lowest-BIC fit retained for subsequent evaluation. Models were evaluated using two complementary criteria: the Bayesian Information Criterion (BIC), which balances statistical goodness-of-fit against model complexity (Schwarz, 1978), and transition-connectivity analysis, which assesses the structural coherence of the inferred state-transition network. Connectivity was evaluated from the directed graph of off-diagonal state transitions with probabilities ⩾ 0.02. Self-transitions were excluded because they describe state persistence rather than communication between behavioural states. Models were considered structurally admissible when the transition graph was strongly connected, contained no isolated state, each hidden state accounted for at least 1% of observations, and the mean maximum posterior probability within every state was ⩾ 0.50. The inferred hidden states were subsequently characterised according to their behavioural properties and transition dynamics. The selected HMM was then implemented within the ABCA framework as an alternative representation of local behavioural dynamics.

### Statistical assumptions

Because imaging was performed in a single focal plane, trajectories were frequently truncated as zoospores left the imaging volume. We therefore assumed that, within each experimental acquisition, local swimming statistics were approximately stationary and that the resulting trajectory fragments provided representative samples of the underlying locomotor process. Individual trajectory fragments were treated as independent realisations conditional on the experimental acquisition. These assumptions underlie both empirical behavioural inference and hidden Markov modelling.

### Simulation framework

#### ABCA software architecture

We developed Agent-Based Cellular Automata (ABCA), an open-source software framework for implementing, simulating, visualising and analysing classical and agent-based cellular automata. ABCA was implemented in OCaml 5.3.0 (ocaml.org) (Leroy *et al*., 2025) using a modular plugin architecture that separates the core simulation engine from model-specific rules and parameters. This design allows biological models to be developed independently while reusing common components for simulation control, visualisation, data export and analysis. The software is built with Dune 3.17.2 (github.com/ocaml/dune) and is publicly available on GitHub (github.com/EEvangelisti/ABCA), facilitating reproducible installation, extension and reuse. Although ABCA is applied here to zoospore motility, its architecture was designed to support the modelling of other biological systems in which individual entities evolve according to local rules.

#### Empirical ABCA model

To reproduce experimentally observed zoospore motility, we implemented an empirical data-driven model as a dedicated ABCA plugin. Model parameters were imported directly from the calibration pipeline. These included state occupancy probabilities and transition probabilities, empirical quantile functions describing swimming-speed and turning-angle distributions, directional turning probabilities, and the transition matrix *A* and innovation covariance matrix *Q* defining the latent bivariate VAR(1) dynamics, together with its stationary covariance matrix *R* (Lutkepohl, 2005). During initialisation, agents were assigned behavioural states according to the experimentally inferred occupancy probabilities. Latent speed and turning variables were generated to reproduce the stationary covariance structure of the calibrated VAR(1) model, while initial headings were distributed uniformly across the full angular range. At each simulation step, the behavioural state of every agent was updated according to the empirical transition matrix, after which the latent variables evolved according to:

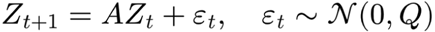

with Gaussian innovations generated using a Cholesky factorisation of *Q*. The updated latent variables were converted into probability ranks through the standard normal cumulative distribution function and mapped onto the corresponding empirical swimming-speed and turning-angle distributions. Swimming speed was additionally constrained by an experimentally calibrated acceleration limit, while turning direction was sampled according to the observed clockwise or counterclockwise probabilities. Internal consistency checks were performed when the calibrated VAR(1) parameters were imported, including verification that the covariance matrices were valid and that the stationarity identity *R* = *ARA*^⊤^ + *Q* was satisfied.

#### HMM-driven ABCA model

As an alternative representation of zoospore behaviour, we implemented a second ABCA plugin driven by the two-state HMM inferred from the experimental trajectories. Each simulated agent carried a latent locomotor state corresponding directly to one of the statistical states identified by the HMM. The plugin was parameterised without re-estimation by importing the initial-state probabilities, state-transition matrix, state-specific empirical distributions of swimming speed and absolute turning angle, and state-specific probabilities of positive versus negative turns. At initialisation, the hidden state of each agent was sampled from the inferred initial-state distribution, while the initial heading was sampled uniformly over 0-360°. At each simulation step, the current hidden state generated one movement observation by independently sampling swimming speed and turning angle from the corresponding state-specific empirical distributions, after which the next hidden state was sampled according to the transition matrix, following the standard HMM generative process (Rabiner, 1989). Unlike the empirical ABCA model, which represents temporal dependence through a continuous VAR(1) process, the HMM-driven model captures temporal organisation exclusively through transitions between latent behavioural states. It therefore provided an alternative translation of experimentally inferred behavioural organisation into agent-level movement rules.

### Simulations and resampling

The empirical and HMM-driven ABCA models were each evaluated using 100 independent simulations, each comprising 2,500 agents followed for 100 time steps. Simulated trajectories were exported in XML format and subsequently resampled to reproduce the empirical trajectory-length distribution. This resampling was necessary because experimental trajectories are frequently truncated when swimming zoospores leave the microscope focal plane, whereas simulated trajectories all have identical duration. The empirical trajectory-length distribution was used directly as the sampling target. For each empirical trajectory length, one simulated trajectory of sufficient length was selected uniformly at random, and a contiguous segment of the required length was extracted from a uniformly sampled starting position. Time indices were then reset so that every resampled trajectory started at *t* = 0, while preserving the original spatial coordinates and local dynamics. This procedure reproduced the empirical trajectory-length distribution while sampling uniformly across eligible simulated trajectories and admissible trajectory segments. The resampled datasets were subsequently analysed using exactly the same pipeline as the experimental data, allowing direct comparison of all trajectory descriptors.

For simulations performed in crowded environments, the obstacle geometry was imported from binary masks generated from the corresponding experimental images by Otsu thresholding. The number of simulated zoospores was fixed to 1,246 agents, corresponding to the mean number of experimentally observed zoospores per frame. Each collision rule was evaluated using 100 independent simulations. Simulations were run for 200 time steps, with the first 60 steps discarded as a burn-in period to remove artefacts arising from random initialisation. The remaining simulated trajectories were then resampled according to the experimental trajectory-length distribution and analysed using the same automated pipeline as the experimental bead datasets.

### Model evaluation

Model performance was evaluated by comparing emergent properties of simulated trajectories with those measured experimentally. Each of the 100 stochastic simulations generated by each model was evaluated independently after resampling to the empirical trajectory-length distribution. Agreement between simulations and experimental observations was assessed using complementary trajectory-level descriptors. Net-displacement distributions were compared using the first-order Wasserstein distance (Ramdas *et al*., 2017), the two-sample Kolmogorov-Smirnov (KS) statistic (Massey, 1951), and the absolute median error **(Table 3)**.

**Table 3.** Quantitative comparison of simulated and experimental zoospore trajectories. Net-displacement distributions were compared using the first-order Wasserstein distance, the Kolmogorov-Smirnov (KS) statistic, and the absolute median error. Mean squared displacement (MSD) curves were compared using the root mean squared error (RMSE) and the mean absolute error (MAE). Values are reported as the median together with the 2.5th-97.5th percentile interval across the 100 simulations. Lower values indicate closer agreement with the experimental data.

| Validation metric | Definition |
| --- | --- |
| First-order Wasserstein distance | $W_1(F_{\text{exp}}, F_{\text{sim}}) = \int_{-\infty}^{+\infty} \vert F_{\text{exp}}(x) - F_{\text{sim}}(x)\vert dx$ |
| Kolmogorov-Smirnov (KS) statistic | $D_{\text{KS}} = \sup_x \vert F_{\text{exp}}(x) - F_{\text{sim}}(x)\vert $ |
| Absolute median error | $E_{\text{med}} = \vert \tilde{x}_{\text{sim}} - \tilde{x}_{\text{exp}}\vert $ |
| Root mean squared error (RMSE) | $\text{RMSE} = \sqrt{\frac{1}{L} \sum_{i=1}^L (M_{\text{sim},i} - M_{\text{exp},i})^2}$ |
| Mean absolute error (MAE) | $\text{MAE} = \frac{1}{L} \sum_{i=1}^L \vert M_{\text{sim},i} - M_{\text{exp},i}\vert $ |

Because the large number of trajectories makes hypothesis-test P-values highly sensitive to minor discrepancies, model comparisons relied on the KS statistic itself rather than on statistical significance. Agreement between experimental and simulated MSD curves was evaluated over their common temporal-lag range. When necessary, simulated MSD values were linearly interpolated to the experimental lag values without extrapolation. Curve discrepancies were quantified using the root mean squared error (RMSE) and the mean absolute error (MAE) **(Table 3)**. Lower values indicate closer agreement between simulated and experimental trajectories for all validation metrics. Each metric was calculated separately for every simulation, and model performance was summarised by the median and 2.5th-97.5th percentile interval across the 100 simulations.

## Results

Large-scale reconstruction of *P. nicotianae* zoospore trajectories.

To quantitatively characterise *Phytophthora* zoospore swimming behaviour, we performed time-lapse imaging of an entire *P. nicotianae* zoospore suspension droplet during exploratory swimming in a liquid environment devoid of external chemical or hydrodynamic cues **(Figure 1A)**. Automated trajectory reconstruction using TrackMate yielded 65,023 individual trajectories, with a median duration of 31 consecutive frames, corresponding to 2.17 s of motion **(Figures 1B and S1A)**. The upper tail of the trajectory-length distribution contained a small proportion of exceptionally long trajectories, which could disproportionately include tracking artefacts or persistent non-biological objects. Subsequent quantitative analyses were therefore performed on a filtered dataset retaining trajectories whose lengths were below the 90th percentile. This filtering step retained 58,696 trajectories, with a median duration of 28 consecutive frames, corresponding to 1.96 s of motion **(Figures 1C and S1B)**. When translated to a common origin, the reconstructed trajectories illustrated the broad diversity of individual swimming patterns across trajectory-length deciles, without revealing a pronounced directional bias **(Figures 1C and S2A-J)**. We then assessed whether instantaneous swimming headings were uniformly distributed throughout the experiment. Analysis of 1,795,669 headings revealed a high degree of directional isotropy, with a resultant vector length of R = 0.053 **(Figure 1D)**. This low value indicates the absence of a strong preferred swimming direction at the population level, although headings around 90° were slightly under-represented. Collectively, these results establish a large-scale experimental trajectory dataset suitable for quantitative analysis and data-driven modelling of zoospore motility.

**Figure 1.**
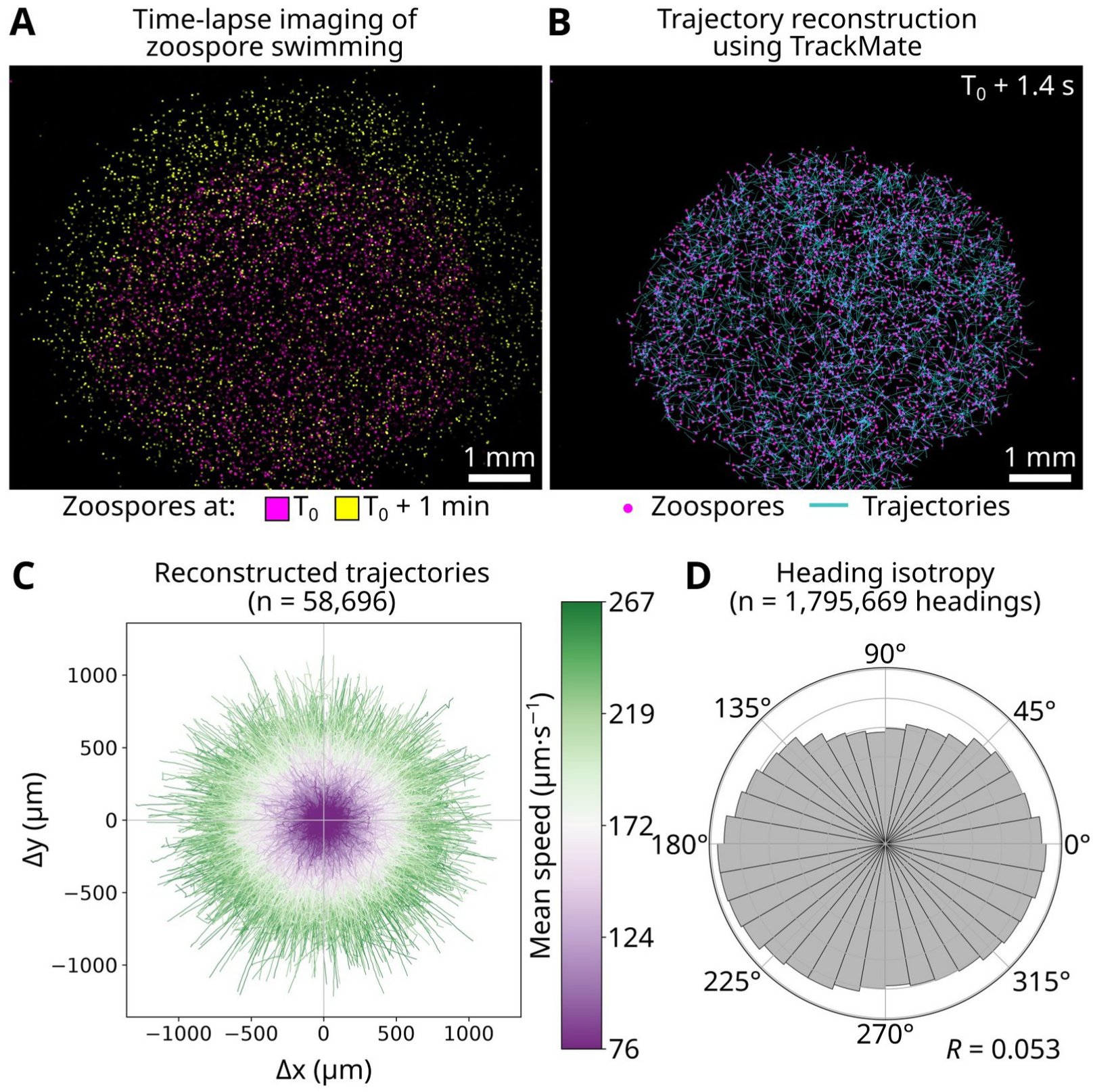
Large-scale reconstruction of *Phytophthora nicotianae* zoospore trajectories. **(A)** Representative images of a *P. nicotianae* zoospore suspension immediately after deposition (T0, magenta) and after 1 min of exploratory swimming (T0 + 1 min, yellow). Zoospores progressively disperse throughout the droplet while remaining evenly distributed in the absence of external cues. **(B)** Automated reconstruction of zoospore trajectories from time-lapse microscopy using TrackMate. Individual detections are shown in magenta and reconstructed trajectories in cyan. **(C)** Overview of the reconstructed trajectory dataset after quality filtering (58,696 trajectories). Each trajectory was translated to the origin to visualise the overall spatial distribution and coloured according to its mean swimming speed. **(D)** Polar histogram of instantaneous swimming headings calculated from all reconstructed trajectories (1,795,669 headings). Each heading corresponds to the orientation of an individual displacement step. The nearly uniform angular distribution (resultant vector length, *R* = 0.053) indicates that zoospore swimming is globally isotropic under these experimental conditions. Scale bars, 1 mm.

### Global trajectory descriptors capture emergent properties of zoospore motility

Next, we analysed the reconstructed trajectories using global trajectory descriptors characterising zoospore exploratory behaviour **(Figure 2)**. The distribution of trajectory straightness was strongly weighted toward high values, with a median of 0.795, indicating that the net displacement of a typical trajectory represented nearly 80% of its total path length **(Figure 2A)**. Thus, zoospore trajectories were predominantly persistent and relatively direct over the observation period, although marked heterogeneity was evident across the population. Straightness nevertheless decreased progressively across trajectory-length deciles, indicating that longer trajectories accumulated greater deviations from a direct path **(Figure S2K)**. We then quantified the net displacement of individual trajectories. The distribution was right-skewed, with a median displacement of 246 µm, while a small fraction of trajectories extended over distances approaching or exceeding 1 mm **(Figure 2B)**. Because trajectories were truncated when zoospores left the field of view or focal plane, these values describe displacement over the reconstructed portion of each swimming path rather than the complete excursion of the cell. The broad displacement distribution was consistent with the progressive outward expansion of the zoospore population observed during time-lapse imaging **(Figure 1A)**. Finally, we quantified the mean squared displacement (MSD), which describes the average squared displacement as a function of lag time. The MSD increased continuously throughout the observation window, reaching approximately 8×10⁴ µm² after 1.75 s **(Figure 2C)**. The progressive widening of the interquartile range further revealed substantial heterogeneity in exploratory behaviour among individual zoospores. Collectively, these global descriptors capture emergent properties of zoospore motility and provide quantitative benchmarks for evaluating whether locally inferred behavioural rules are sufficient to reproduce population-scale exploratory behaviour.

**Figure 2.**
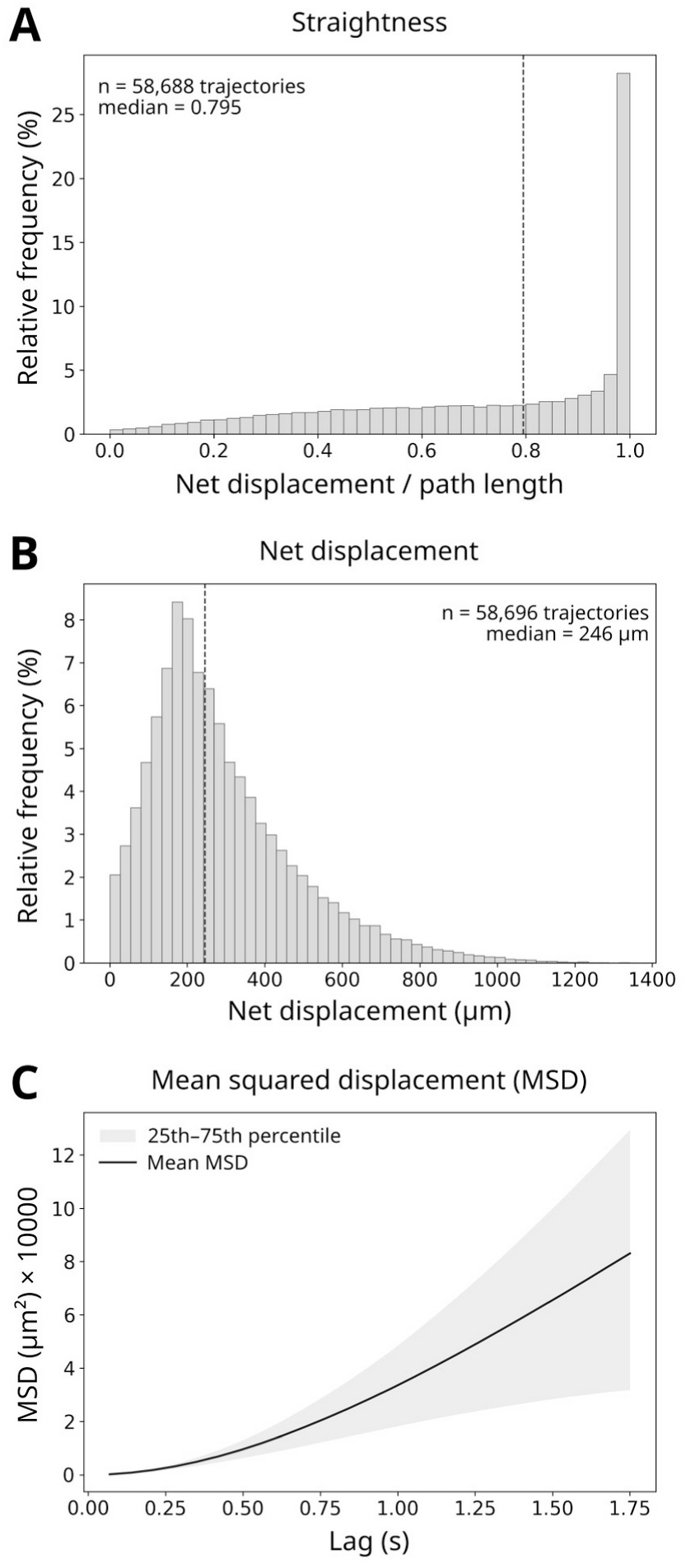
Global trajectory descriptors capture emergent properties of *Phytophthora nicotianae* zoospore motility. **(A)** Distribution of straightness across reconstructed trajectories, defined as the ratio of net displacement to total path length. The dashed vertical line indicates the median straightness. **(B)** Distribution of net displacement for individual reconstructed trajectories. The dashed vertical line indicates the median net displacement. **(C)** Mean squared displacement (MSD) as a function of lag time. The solid line represents the mean MSD, while the shaded area indicates the interquartile range (25th-75th percentile) across trajectories. The continuous increase in MSD with lag time and the widening interquartile range reflect increasing spatial dispersion and trajectory-to-trajectory variability.

### Instantaneous swimming descriptors characterise local zoospore motility

We next examined instantaneous descriptors characterising successive steps of zoospore swimming trajectories **(Figure 3)**. The distribution of swimming speeds was right-skewed, with a median of 211 µm·s^−1^ across 1,795,669 trajectory steps **(Figure 3A)**. Instantaneous speeds above 300 µm·s^−1^ were uncommon (50,102 observations; 2.8%), indicating that very rapid swimming represented only a small fraction of recorded movements, some of which may reflect trajectory-reconstruction artefacts. These values were consistent with previous quantitative measurements of *P. nicotianae* zoospore motility obtained by high-speed imaging (Tran *et al*., 2022). Absolute acceleration was also strongly right-skewed, with a median of 471 µm·s^−2^ across 1,736,973 trajectory steps **(Figure 3B)**, showing that large accelerations or decelerations occurred relatively infrequently. The distribution of signed turning angles was sharply centred around zero, with a median of −0.0578° **(Figure 3C)**. This near-zero value indicates the absence of an apparent population-level bias towards clockwise or anticlockwise turning, consistent with the overall directional isotropy observed across the complete dataset. Examination of absolute turning angles further showed that most successive changes in heading were small, with a median of 5.98° **(Figure 3D)**. Large reorientation events nevertheless occurred, although they represented only a minor fraction of trajectory steps. Collectively, these local behavioural descriptors indicate that zoospore swimming is dominated by relatively fast, directionally persistent motion, punctuated by less frequent large changes in swimming speed and occasional pronounced reorientation events. This organisation is consistent with previous high-speed analyses showing that *Phytophthora* zoospores alternate between persistent swimming runs and rapid turning episodes (Tran *et al*., 2022; Kasteel *et al*., 2024), and provides the quantitative basis for inferring local behavioural rules.

**Figure 3.**
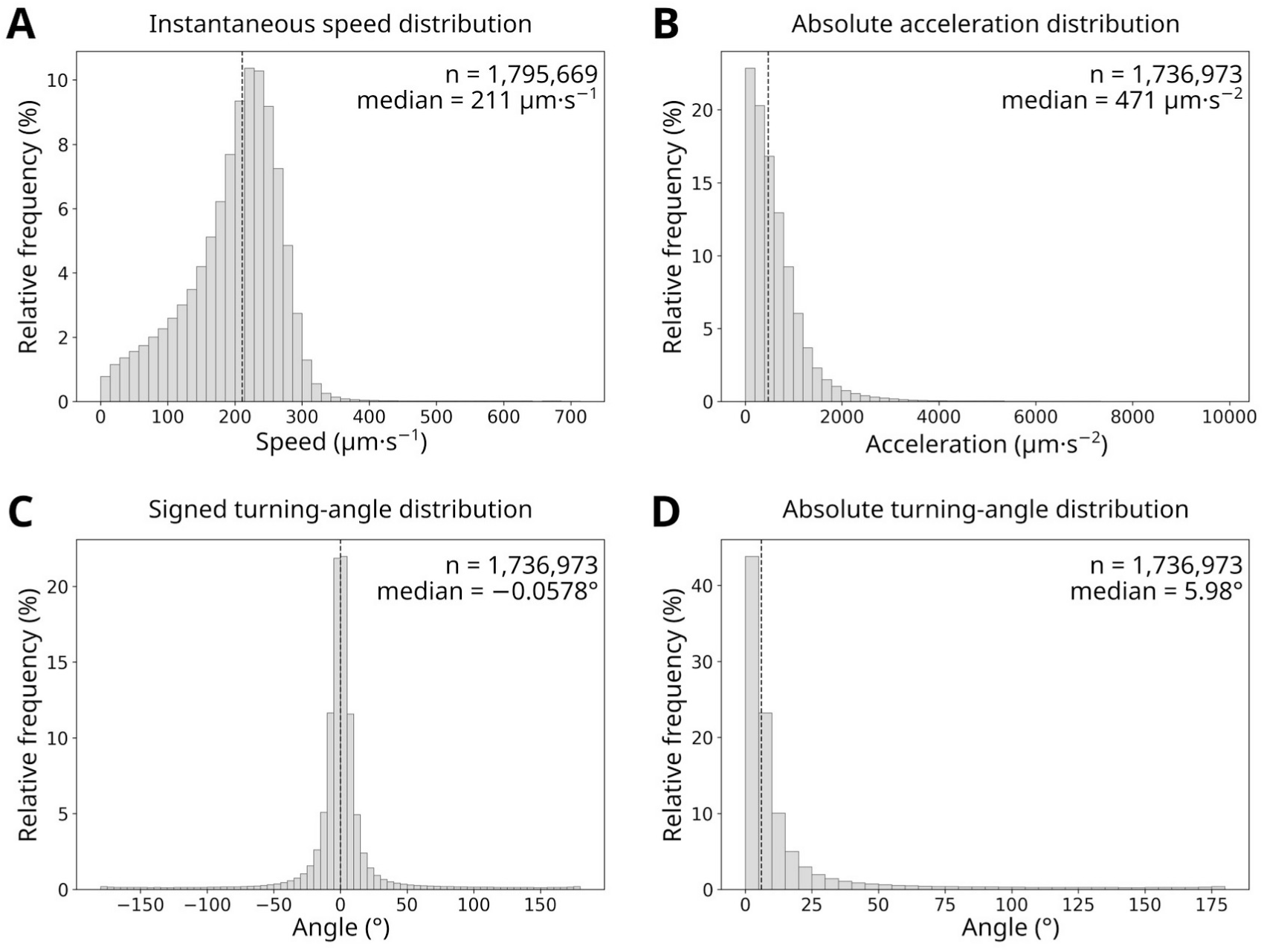
Instantaneous swimming descriptors characterise local zoospore motility. **(A)** Distribution of instantaneous swimming speeds measured across all reconstructed trajectory steps. **(B)** Distribution of absolute acceleration values computed between successive trajectory steps. **(C)** Distribution of signed turning angles computed between successive trajectory segments. The distribution is centred around 0°, indicating the absence of a preferred clockwise or anticlockwise turning direction. **(D)** Distribution of absolute turning angles computed between successive trajectory segments. Dashed vertical lines indicate the median values.

### Local swimming descriptors exhibit temporal memory and cross-variable coupling

We next investigated temporal and cross-variable dependencies among the local motility descriptors **(Figure 4; Figure S3)**, because these dependencies determine whether successive swimming variables can be sampled independently in a behavioural model. We first examined the temporal autocorrelation of swimming speed by calculating, for each trajectory and lag time, the Pearson correlation between instantaneous speeds separated by the corresponding lag **(Figure 4A)**. Speed autocorrelation was modest, reaching a maximum mean value of 0.16 at a lag of 0.14 s, and rapidly approached zero at longer lag times. Nevertheless, trajectory-level autocorrelation coefficients were consistently greater than zero at every lag tested (Wilcoxon signed-rank tests, Benjamini-Hochberg (BH)-adjusted *q* < 0.001 for all lags). Thus, swimming speed exhibited a statistically robust but short-lived and quantitatively limited temporal dependence. We then assessed directional persistence as the mean cosine of the change in heading between positions separated by each lag time **(Figure 4B)**. In marked contrast to swimming speed, directional persistence reached 0.86 at a lag of 0.14 s and declined only gradually, remaining positive on average at the longest lag examined, 1.75 s. Directional persistence also differed significantly from zero across all tested lags (Wilcoxon signed-rank tests, BH-adjusted *q* < 0.001). These results indicate that swimming speed retains only short-lived temporal memory, whereas swimming direction remains persistent over substantially longer timescales.

**Figure 4.**
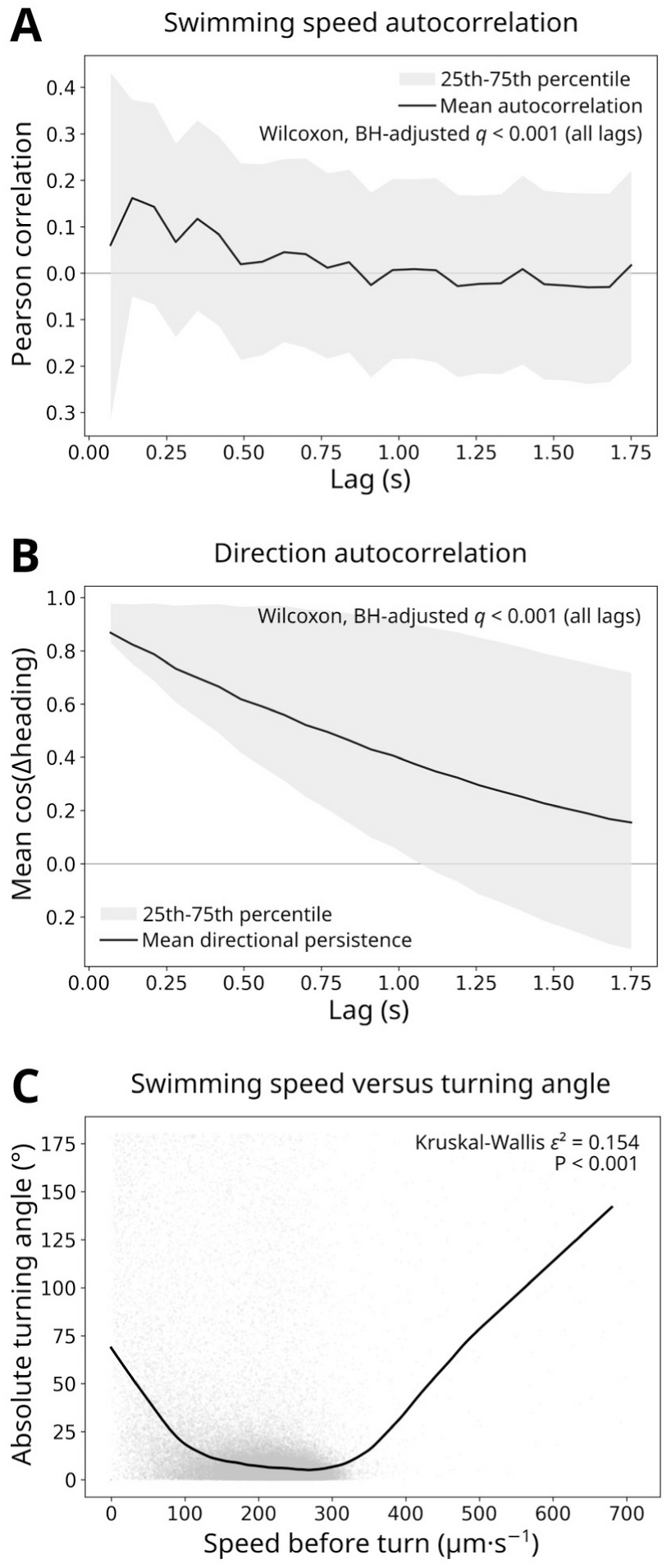
Local swimming descriptors exhibit temporal memory and cross-variable coupling. **(A)** Mean Pearson autocorrelation coefficient of instantaneous swimming speed as a function of lag time. The black line shows the mean autocorrelation calculated independently for each trajectory, and the grey shaded area indicates the interquartile range (25th-75th percentile). **(B)** Directional persistence as a function of lag time, calculated as the mean cosine of the change in heading (Δheading) between positions separated by each lag. The black line indicates the mean directional persistence across trajectories, and the grey shaded area represents the interquartile range (25th-75th percentile). For both panels, Wilcoxon signed-rank tests against zero remained significant for all lag times after Benjamini-Hochberg correction (BH-adjusted *q* < 0.001). **(C)** Relationship between swimming speed immediately preceding each change in heading and the corresponding absolute turning angle. Grey points represent individual observations, and the black curve shows a locally weighted regression (LOESS). Differences in turning-angle distributions across speed deciles were assessed using a Kruskal-Wallis test (P < 0.001, ε² = 0.154).

Finally, we examined the relationship between swimming speed and absolute turning angle **(Figure 4C)**. Turning angles were largest at low swimming speeds and reached their lowest values at intermediate-to-high speeds of approximately 200-300 µm·s^−1^, indicating that higher swimming speeds were generally associated with smaller changes in direction. Turning angles increased again at the extreme upper end of the speed distribution. Because relatively few observations were available in this range **(Figure 3A)**, and some may reflect trajectory-reconstruction artefacts, this apparent increase should be interpreted cautiously. Turning-angle distributions nevertheless differed substantially among speed deciles (Kruskal-Wallis test, P < 0.001, *ε*² = 0.154). A significant relationship was also observed between swimming speed and acceleration **(Figure S3)**. However, because acceleration is calculated from successive speed measurements, this association is expected mathematically and therefore does not constitute independent evidence of biological coupling. Consequently, acceleration was not considered as an independent variable for construction of the empirical behavioural model. Together, these findings demonstrate that local zoospore motility cannot be fully represented by independently sampling empirical speed and turning-angle distributions. Instead, realistic behavioural models must preserve both temporal memory and cross-variable coupling. These statistical dependencies therefore constitute essential constraints for data-driven inference of local behavioural rules.

### A data-driven behavioural model parameterised directly from experimental trajectories

We next investigated whether zoospore swimming behaviour could be reproduced using an agent-based cellular automaton (ABCA). The most parsimonious formulation would represent all observations as arising from a single continuous motility regime. However, the pronounced and non-linear dependence of turning behaviour on swimming speed indicated that local movement statistics varied substantially across the speed distribution **(Figure 4C)**. In particular, low-speed observations were associated with markedly larger directional changes, whereas intermediate-to-high swimming speeds were predominantly associated with persistent, straighter motion. These observations motivated a minimal discrete approximation in which local movement statistics were partitioned into two empirical behavioural regimes, hereafter termed SLOW and FAST. This two-state representation was not intended to assume the existence of two intrinsically distinct biological programmes, but to provide the simplest partition capable of preserving state-dependent speed and turning statistics, their temporal persistence, and transitions between contrasting local swimming conditions.

To initialise the partition objectively, we applied Otsu’s threshold-selection method (Otsu, 1979), which determines the threshold that maximises the between-class variance of the swimming-speed distribution. This yielded an initial threshold of 171 µm·s^−1^. Direct application of this threshold to the experimental trajectories resulted in median SLOW- and FAST-episode durations of 0.07 s and 0.21 s, respectively **(Figure S4)**. Because 0.07 s corresponds to a single acquisition interval in our experimental setup, this frame-wise partition generated frequent state alternations, indicative of threshold-induced flickering rather than stable behavioural transitions.

To stabilise state assignment, we introduced a hysteresis interval around the Otsu threshold and evaluated a range of hysteresis half-widths **(Figures 5A and S5)**. Increasing the hysteresis progressively reduced the proportion of one-step episodes and increased the temporal coherence of both behavioural states. However, wider hysteresis intervals also progressively merged neighbouring episodes and eventually altered overall state occupancy. We therefore retained a hysteresis half-width of 25 µm·s^−1^. This increased the median duration of

**Figure 5.**
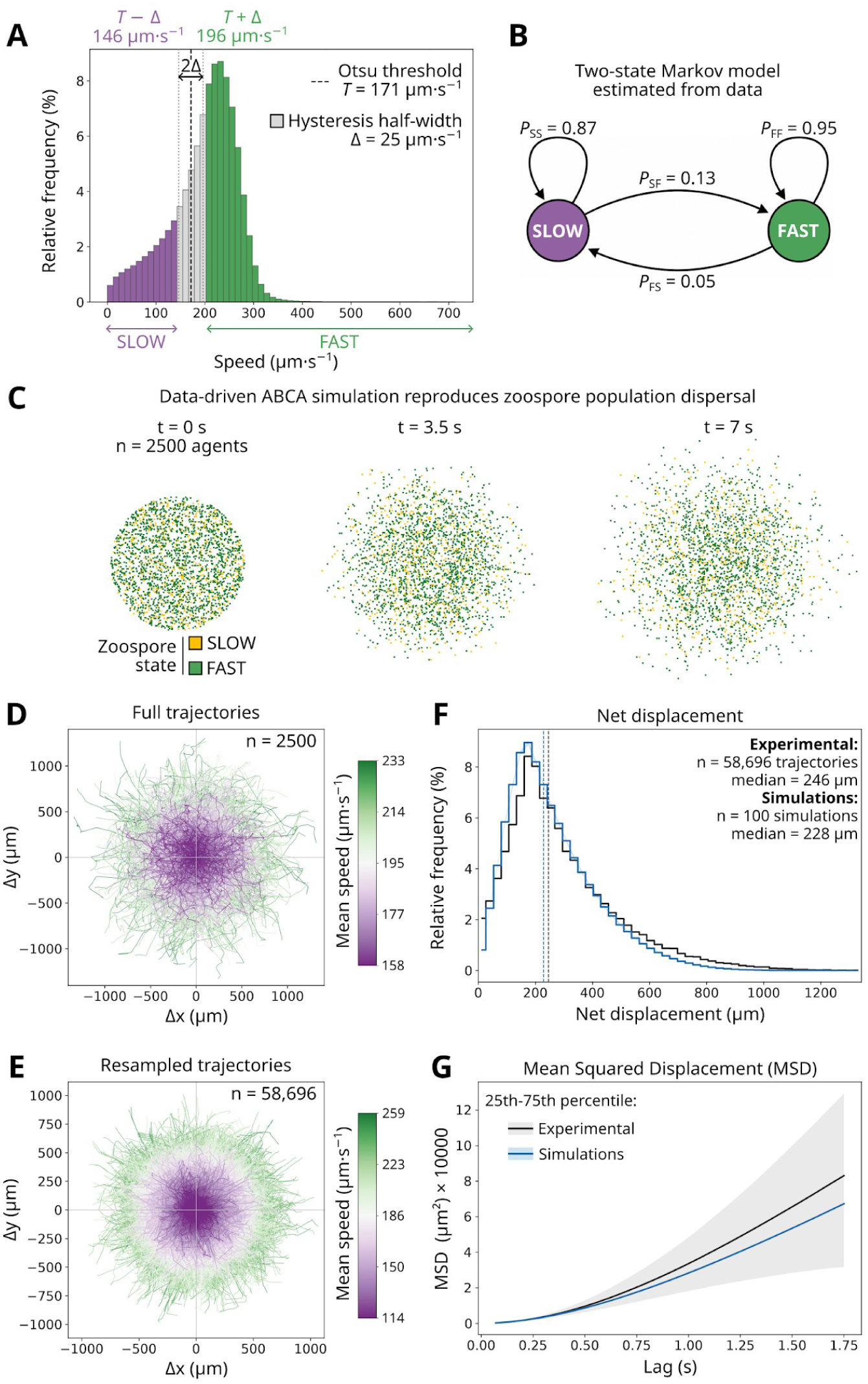
A data-driven empirical ABCA model recapitulates emergent properties of *P. nicotianae* zoospore swimming. **(A)** Instantaneous swimming speeds were partitioned into SLOW and FAST behavioural states using an Otsu threshold (*T* = 171 µm·s⁻¹) and a hysteresis half-width of Δ = 25 µm·s⁻¹. **(B)** State-transition probabilities estimated from the experimental trajectories were used to parameterise a two-state behavioural model. *P*, probability; F, FAST; S, SLOW. **(C)** Representative simulation of 2,500 zoospores showing the progressive expansion of the population over time. Zoospores are coloured according to their behavioural state (SLOW, orange; FAST, green). **(D)** Complete simulated trajectories. **(E)** Simulated trajectories resampled to match the empirical trajectory-length distribution. **(F-G)** Comparison of experimental (grey) and simulated (blue) net-displacement distributions **(F)** and mean squared displacement (MSD) curves **(G)**. Dashed vertical lines indicate median values.

FAST and SLOW episodes to 0.63 s and 0.21 s, respectively, while substantially reducing one-step episodes and preserving the overall balance between behavioural states **(Figure S5)**.

These experimentally inferred behavioural rules were then implemented in the ABCA framework without further parameter adjustment. We performed 100 independent stochastic simulations, each initialised with a different pseudorandom seed and comprising 2,500 agents followed for 100 simulation steps **(Figure S6)**. Simulated trajectories were subsequently resampled to reproduce the experimental trajectory-length distribution before quantitative comparison **(Figure S6A)**. Examples of complete and resampled simulated trajectories are shown in **Figures 5D** and **5E**, respectively, while their organisation across trajectory-length deciles is shown in **Figure S7**. We first verified that the simulations faithfully reproduced the local descriptors used for model calibration. The distributions of instantaneous swimming speed, signed and absolute turning angle, and signed and absolute acceleration closely matched their experimental counterparts across the 100 independent simulations **(Figure S6B-F)**. As expected, variability among simulations was minimal for these local descriptors because each simulation contained a large number of observations and was generated from the same experimentally calibrated stochastic process.

We next evaluated whether these locally inferred behavioural rules were sufficient to reproduce trajectory-scale properties that had not been used during model calibration. The simulated net-displacement distribution closely reproduced the broad shape of the experimental distribution, although its median was moderately reduced, from 246 µm experimentally to approximately 228 µm in the simulations **(Figures 5F)**. Similarly, the simulated MSD followed the characteristic increase observed experimentally but remained progressively lower at longer lag times **(Figures 5G)**. These concordant differences indicate a modest underestimation of long-range exploration rather than a failure to reproduce the overall geometry of zoospore movement. Together, these results show that behavioural rules inferred exclusively from local trajectory statistics capture the principal emergent spatial properties of experimental zoospore trajectories, while modestly underestimating long-range exploration.

### Hidden Markov modelling independently recovers the empirical behavioural organisation

We next investigated whether hidden Markov models (HMMs) fitted directly to the experimental trajectories would independently recover the behavioural organisation inferred by the empirical model. HMMs containing two to seven hidden states were evaluated using the Bayesian information criterion (BIC) together with the transition-connectivity criterion described above **(Figure S8)**. Although increasing the number of hidden states improved model fit, all models containing three or more states introduced at least one short-lived state with a median sojourn duration of a single acquisition interval **(Figure S8)**. We therefore retained the two-state HMM as the most parsimonious model yielding temporally persistent behavioural states and examined it in detail **(Figure 6A, S9)**. The transition structure of the two-state HMM closely resembled that of the empirical SLOW/FAST model. Transition probabilities between hidden states were similar to the experimentally inferred SLOW/FAST transition probabilities, with comparable switching frequencies and state persistence **(Figure 6A)**. Consistent with this correspondence, 81.8% of observations assigned to HMM state 0 belonged to the empirical SLOW state, whereas 93.1% of observations assigned to HMM state 1 belonged to the empirical FAST state **(Figure 6B)**.

**Figure 6.**
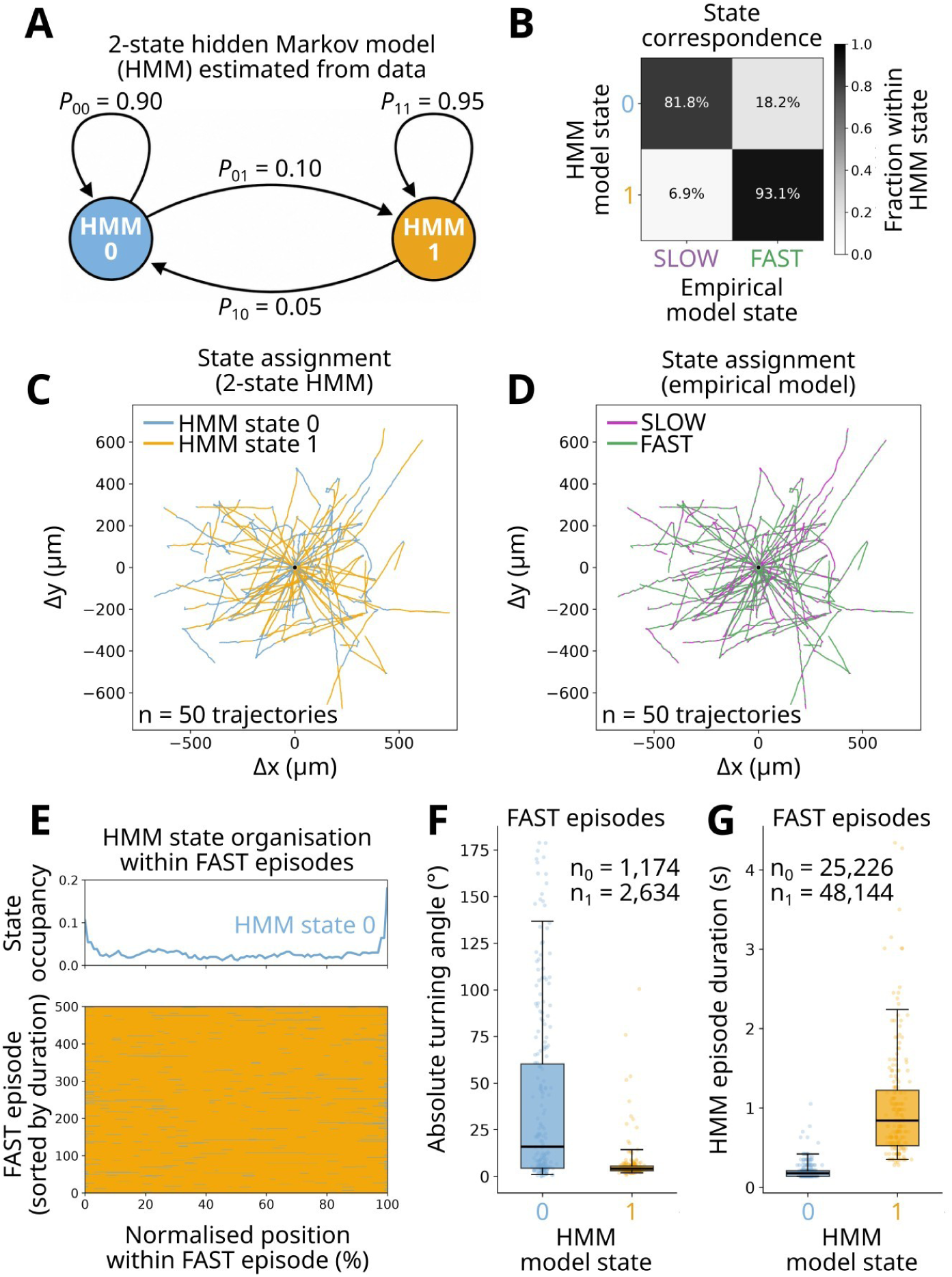
Hidden Markov modelling refines the behavioural interpretation of empirical zoospore swimming states. **(A)** Two-state hidden Markov model (HMM) inferred from experimental trajectory data. Transition probabilities indicate strong persistence of both hidden states. *P*, probability; 0: HMM state 0, 1: HMM state 1. **(B)** Correspondence between HMM states and the empirical SLOW/FAST classification. **(C-D)** Representative centred trajectories coloured according to HMM state assignment **(C)** or the empirical SLOW/FAST classification **(D)**. **(E)** Distribution of HMM states within empirical FAST episodes. The upper panel shows the mean occupancy of HMM state 0 as a function of normalised episode duration; the lower panel shows individual FAST episodes sorted by duration. **(F)** Absolute turning angles of HMM states within empirical FAST episodes. **(G)** Duration of contiguous HMM state episodes within empirical FAST episodes.

Detailed examination of the hidden states further refined their behavioural interpretation. HMM state 0 was characterised by lower swimming speed and larger turning angles than HMM state 1, whereas acceleration contributed comparatively little to state discrimination despite reaching strong statistical significance because of the large sample size **(Figure S9A-C)**. The median sojourn durations of both hidden states exceeded the acquisition interval and closely matched the persistence of the empirical SLOW and FAST states **(Figure S9D, E)**. Together, these observations indicate that the latent states recovered by the HMM largely correspond to the empirical partition, providing independent, unsupervised support for its underlying behavioural organisation.

To determine how the two classifications differed despite their overall correspondence, we mapped both state assignments onto the same experimental trajectories **(Figures 6C, D)**. Visual inspection showed that HMM state 0 was preferentially associated with pronounced changes in swimming direction, whereas HMM state 1 predominantly occupied long, directionally persistent trajectory segments **(Figure 6C)**. By contrast, empirical SLOW and FAST labels were more diffusely distributed along trajectories and were less specifically associated with individual reorientation events **(Figure 6D)**. Thus, although the two approaches recovered closely related behavioural organisations, the HMM was not simply reproducing a speed-based partition, but incorporated additional information related to directional dynamics. To investigate this difference further, we examined the organisation of HMM states within trajectory segments classified as FAST by the empirical model **(Figures 6E-G)**. As expected from the overall state correspondence, empirical FAST episodes were predominantly occupied by HMM state 1 **(Figure 6E)**. However, a reproducible minority of HMM state 0 observations occurred within these episodes, with a clear enrichment near their beginning and end. These embedded HMM state 0 events were associated with substantially larger turning angles than HMM state 1 observations and had markedly shorter durations **(Figures 6F,G)**.

Together, these analyses show that the two independently developed approaches recover closely related principal behavioural organisations of zoospore swimming. However, the HMM further reveals that empirical FAST episodes are not behaviourally homogeneous. Rather, they consist primarily of sustained, directionally persistent swimming interspersed with brief reorientation events. The hidden states therefore integrate swimming speed, directional persistence and turning dynamics, refining the empirical SLOW/FAST description while independently supporting its overall behavioural organisation.

### Empirical and HMM-derived behavioural rules capture complementary features of zoospore dispersal

Since the two-state HMM independently recovered a closely related behavioural organisation, we next investigated whether it could also serve as a generative model of zoospore dispersal. The inferred HMM parameters were implemented directly within the ABCA framework, and 100 independent simulations were performed. A representative simulation illustrates the progressive expansion of a population of 2500 zoospores over time **(Figure 7A)**. Complete simulated trajectories **(Figure 7B)** were subsequently resampled to reproduce the empirical trajectory-length distribution before quantitative comparison **(Figure 7C, S10)**. The resulting simulations reproduced the main emergent properties of experimental dispersal. The net-displacement distribution closely matched that measured experimentally **(Figure 7D)**, including a nearly identical median displacement (278 µm versus 246 µm experimentally). Likewise, simulated MSD closely followed the experimental dynamics throughout the observation period **(Figure 7E)**, indicating that the HMM-derived behavioural states were sufficient to reproduce the principal population-scale dispersal properties. Quantitative comparison revealed complementary strengths of the two generative formulations **(Table 4)**. For net displacement, neither model consistently outperformed the other across all metrics. The HMM-driven model produced a slightly lower Wasserstein distance than the empirical SLOW/FAST model (30.54 versus 32.54 µm, respectively), whereas the empirical model provided a lower Kolmogorov–Smirnov statistic (0.0576 versus 0.0741, respectively) and a smaller median displacement error (17.91 versus 31.58 µm, respectively). By contrast, the HMM-driven model consistently reproduced MSD dynamics more accurately, reducing RMSE from 7596.53 to 5301.15 µm² and MAE from 5633.50 to 3811.80 µm², corresponding to reductions of approximately 30% and 32%, respectively.

**Figure 7.**
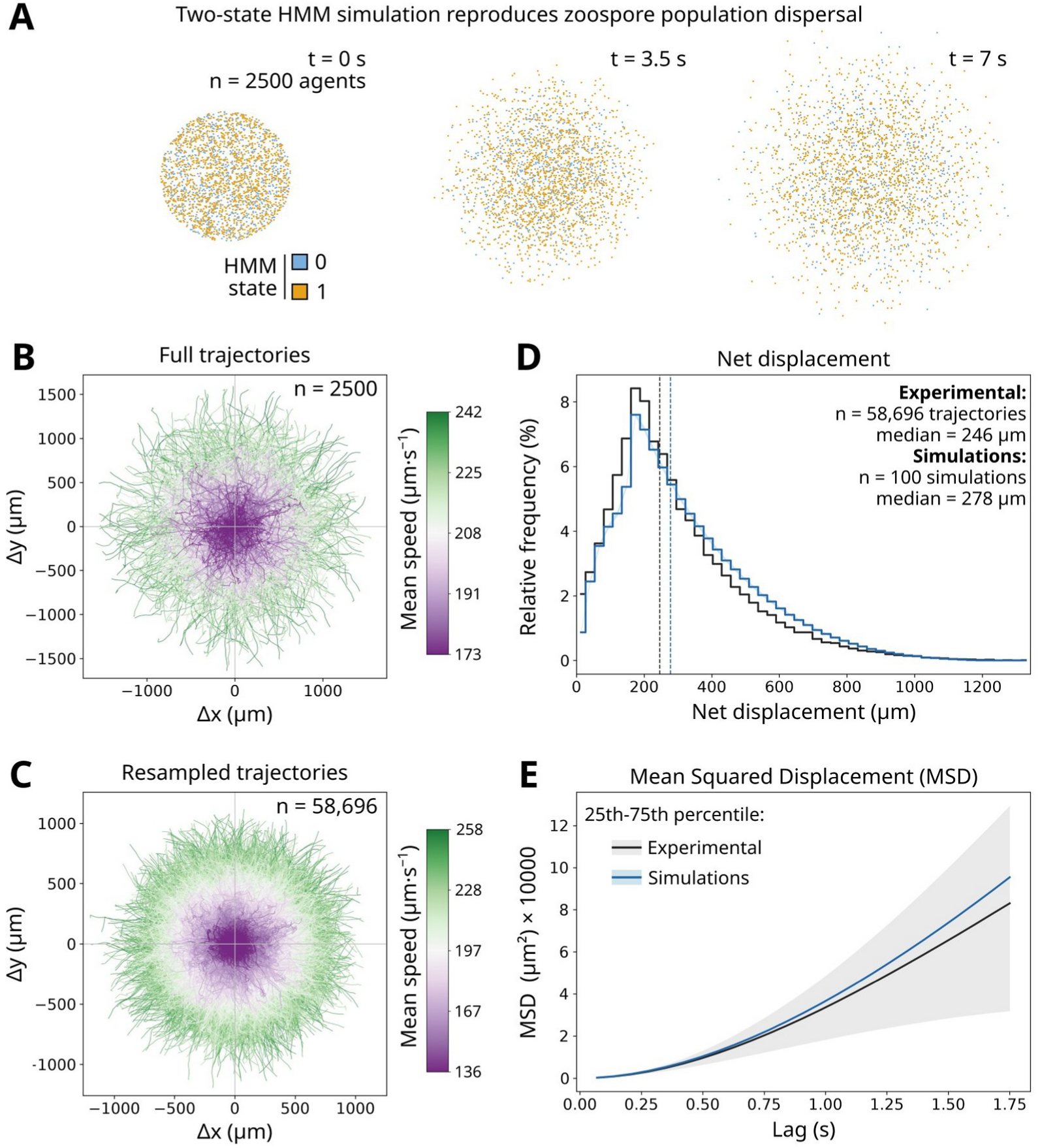
A two-state HMM-driven ABCA model reproduces emergent properties of *P. nicotianae* zoospore dispersal. **(A)** Representative simulation of 2,500 zoospores showing the progressive expansion of the population over 7 s. Agents are coloured according to their inferred HMM state (state 0, blue; state 1, orange). **(B)** Complete simulated trajectories coloured according to mean swimming speed. **(C)** Simulated trajectories resampled to match the empirical trajectory-length distribution. **(D-E)** Comparison of experimental (black) and simulated (blue) net-displacement distributions **(D)** and mean squared displacement (MSD) curves **(E)**. Vertical dashed lines indicate median values. Shaded areas represent the 25th-75th percentile interval across simulations **(D)** or trajectories **(E)**.

**Table 4.** Quantitative comparison of the empirical SLOW/FAST and two-state HMM-driven ABCA models. Agreement between simulated and experimental data was evaluated using complementary global descriptors of zoospore dispersal. Net-displacement distributions were compared using the first-order Wasserstein distance, the two-sample Kolmogorov-Smirnov (KS) statistic, and the absolute median error. Mean squared displacement (MSD) curves were compared using the root mean squared error (RMSE) and mean absolute error (MAE). Lower values indicate better agreement with the experimental data. Reported values correspond to the median across 100 independent simulations, with the 95% interval shown in brackets.

| Descriptor | SLOW/FAST model | Two-state HMM | Best model |
| --- | --- | --- | --- |
| Net displacement Wasserstein | 32.54 [30.50-34.35] | 30.54 [27.43-34.57] | HMM: 6% |
| Net displacement KS | 0.0576 [0.0521-0.0622] | 0.0741 [0.0683-0.0813] | S/F: 22% |
| Median displacement error ($\mu\text{m}$) | 17.91 [15.76–20.64] | 31.58 [28.97–35.24] | S/F: 43% |
| MSD RMSE ($\mu\text{m}^2$) | 7596.5 [7149.9-8065.6] | 5301.1 [4599.2-5974.4] | HMM: 30% |
| MSD MAE ($\mu\text{m}^2$) | 5633.5 [5319.4-5977.9] | 3811.8 [3287.2-4292.3] | HMM: 32% |

Thus, incorporation of the HMM-derived behavioural organisation did not uniformly improve all emergent descriptors. Rather, the two-state HMM improved the temporal dynamics of spatial exploration while the simpler empirical SLOW/FAST formulation remained equally or more accurate for several properties of the net-displacement distribution. These results indicate that the HMM-derived representation better captures specific aspects of dispersal dynamics, particularly the temporal evolution of spatial exploration, while the additional complexity of this representation is not required to recover the overall spatial outcome.

### Experimentally inferred swimming rules predict zoospore dispersal in a novel obstacle environment

As a stringent predictive test of the empirical behavioural model, we next examined whether movement rules inferred exclusively from unobstructed swimming were sufficient to reproduce zoospore behaviour in a physically constrained environment. To this end, we recorded the swimming of *P. palmivora* zoospores in a dense suspension of 10 µm polystyrene microspheres, generating a heterogeneous environment in which swimmers repeatedly encountered immobile obstacles **(Figure 8A)**. The spatial distribution of microspheres was extracted directly from the experimental images and reproduced within the ABCA framework, ensuring that simulated zoospores experienced the experimentally observed obstacle geometry **(Figure 8B)**. Importantly, none of these experiments or parameters contributed to the calibration of the empirical swimming model, providing an independent test of the model beyond the conditions used for calibration.

**Figure 8.**
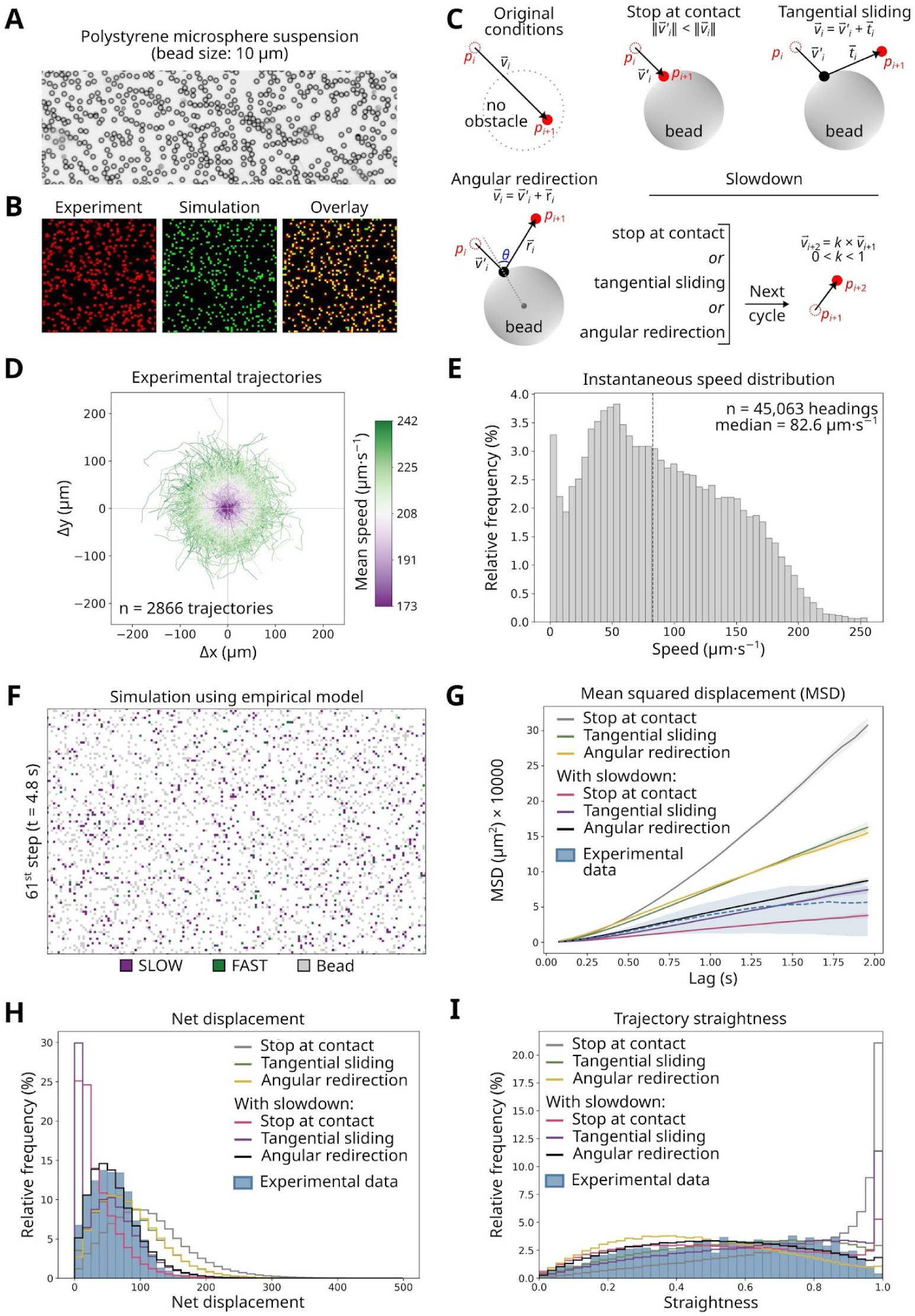
Predictive validation of the empirical zoospore model in a bead-filled environment using alternative collision rules. **(A)** Bright-field image of the experimental suspension of 10 µm polystyrene microspheres. **(B)** Comparison between the experimental bead distribution (red), the corresponding simulation (green), and their overlay (yellow). **(C)** Collision rules evaluated in the model. ‘Stop at contact’ terminates the current displacement at the collision point. ‘Tangential sliding’ projects the remaining displacement onto the local tangent of the bead surface. ‘Angular redirection’ reorients the remaining displacement by a random angle relative to the surface normal. ‘Slowdown’ supplements each geometric collision rule with a transient post-collision reduction in swimming speed. **(D)** Experimental trajectories of *P. palmivora* swimming in the bead suspension, coloured according to mean swimming speed. The mean number of simultaneously tracked zoospores was 1246. **(E)** Experimental distribution of instantaneous swimming speeds. **(F)** Representative simulation using the empirical swimming model and the experimentally derived bead map. **(G-I)** Comparison of alternative collision rules using mean squared displacement **(G)**, net displacement **(H)**, and trajectory straightness **(I)**. All simulations used the same empirical swimming model, identical bead geometry and identical simulation settings; only the collision rule differed (100 simulations per collision rule). Dashed lines and blue histograms indicate the experimental data.

Because the empirical SLOW/FAST model does not specify how zoospores interact with physical obstacles, we evaluated four elementary collision rules together with their combinations **(Figure 8C)**. These rules represented alternative hypotheses describing the immediate response of a zoospore upon contacting a bead: termination of the current displacement (‘stop at contact’), projection of the remaining displacement along the bead surface (‘tangential sliding’), angular reorientation away from the obstacle (‘angular redirection’), each evaluated either alone or in combination with a transient reduction of swimming speed (‘slowdown’). All simulations used the same empirical swimming model, identical bead geometry and identical simulation settings; only the collision rule differed between simulations. Each collision hypothesis was evaluated across 100 independent simulations. To minimise crowding effects arising from zoospore-zoospore interactions, the number of simulated agents was matched to the mean number of experimentally observed zoospores per frame (1,246 agents).

Experimental trajectories revealed that zoospores remained motile within the bead-filled environment while exhibiting substantially reduced dispersal compared with unobstructed swimming **(Figure 8D-F)**. Comparison of the simulated and experimental emergent descriptors discriminated between the competing collision hypotheses **(Figure 8G-I)**. None of the collision rules that preserved swimming speed (‘stop at contact’, ‘tangential sliding’ or ‘angular redirection’) reproduced the experimental dispersal, indicating that the purely geometric collision responses tested here were insufficient to account for zoospore behaviour in the bead-filled environment. By contrast, incorporating a transient post-collision slowdown consistently improved agreement with the experimental data, indicating that, among the collision responses tested, a transient reduction in swimming speed was required to account for the experimental dispersal. Slowdown combined with ‘stop at contact’ underestimated exploratory behaviour and generated trajectories that were less dispersed than those observed experimentally. The best overall agreement was obtained by combining slowdown with ‘angular redirection’, which simultaneously reproduced the experimental mean squared displacement together with the distributions of net displacement and trajectory straightness. Together, these results identify transient post-collision slowdown as a key effective response required to reproduce zoospore dispersal in the bead-filled environment, while additional directional reorientation further improves quantitative agreement.

## Discussion

We developed a data-driven framework to infer and simulate the exploratory swimming behaviour of *Phytophthora* zoospores from a large experimental dataset. Because long-distance three-dimensional tracking of freely swimming zoospores remains technically challenging, we analysed numerous short two-dimensional trajectory fragments. Their large number nevertheless allowed us to characterise motility at two complementary scales: local descriptors, including instantaneous speed, turning, acceleration, behavioural temporal persistence and speed-turn coupling, were used to infer movement rules, whereas global properties such as net displacement and mean squared displacement provided trajectory-scale benchmarks for model evaluation.

Two complementary inference strategies converged on a similar coarse-grained organisation of zoospore swimming. The empirical model separated trajectories into SLOW and FAST regimes, consistent with previous observations that zoospore swimming alternates between rapid directional motion and reorientation events (Tran *et al*., 2022). An independently inferred hidden Markov model nevertheless recovered two latent states defined jointly by swimming speed, turning dynamics and directional persistence. Multivariate definitions of motility states have been reported across diverse systems, including collective cell migration (Allen *et al*., 2020), fibroblast migration (Davis *et al*., 2026) and intracellular particle motion (Röding *et al*., 2014). The agreement between these approaches suggests that the two-regime organisation captured by the empirical model reflects a reproducible structure in the experimental data rather than an arbitrary discretisation imposed by the modeller. The HMM further refined this description by showing that behavioural states could not be reduced to speed alone.

This organisation also differs from a simple run-and-tumble representation. Classical run-and-tumble descriptions separate relatively persistent directional runs from discrete reorientation events (Wadhwa & Berg, 2022). By contrast, the states inferred here represent broader dynamical regimes in which speed, turning amplitude and directional persistence covary, while retaining temporal memory within each state. The HMM therefore provides a more nuanced generative description of exploratory swimming than a simple alternation between directional motion and stochastic reorientation. At the same time, its convergence with the empirical SLOW/FAST model highlights the value of combining interpretable behavioural classifications with latent-state inference to test the robustness of data-driven movement rules.

The central challenge, however, is not only whether inferred local rules can reproduce the data from which they were obtained, but whether they generalise when conditions change. We therefore challenged swimming rules inferred from freely swimming *P. nicotianae* in an independent experiment using *P. palmivora* navigating an obstacle-filled environment. This test went beyond validation on additional trajectories acquired under the same conditions in two respects: both the biological system and the physical environment were changed. Importantly, the underlying swimming rules were not recalibrated. Instead, environmental complexity was introduced through additional rules governing interactions with obstacles.

The need for an additional collision rule does not imply failure of the inferred swimming model, because intrinsic agent behaviour and interactions with the surrounding environment constitute distinct components of an agent-based system (Murphy *et al*., 2020; Griesemer & Sindi, 2022). The former describes unconstrained agent behaviour, whereas the latter specifies how that behaviour is modified by environmental encounters. Purely geometric collision responses were insufficient to reproduce the experimentally observed reduction in dispersal, whereas rules incorporating transient post-collision slowdown substantially improved model agreement. Predictive failure was therefore informative: it identified an additional effective response required specifically when zoospores encounter physical obstacles. The model thus retained its intrinsic behavioural organisation while revealing which new rule had to be introduced when environmental complexity changed.

Such effective rules should not be interpreted as direct descriptions of the molecular or biomechanical mechanisms that generate them. Agent-based models can demonstrate that a given set of local rules is sufficient to reproduce an emergent behaviour without necessarily establishing that these rules correspond one-to-one with underlying biological mechanisms (Glen *et al*., 2019). In the present framework, SLOW and FAST regimes or HMM states should therefore not be interpreted as discrete flagellar or molecular programmes. Rather, they provide compact statistical descriptions of the observable consequences of those processes at the trajectory level. This abstraction is deliberate: the aim of the model is not to reconstruct every local molecular event, but to determine whether experimentally inferred behavioural dynamics provide a sufficient effective description to predict population-level dispersal.

This distinction also clarifies the relationship between our framework and biophysical studies of zoospore propulsion. *Phytophthora* swimming depends on the coordinated action of two structurally distinct flagella, and high-speed imaging combined with biophysical modelling has shown that changes in their coordination can generate both rapid directional swimming and sharp reorientation events (Tran *et al*., 2022). Such approaches explain how local movements may arise mechanistically, whereas the present framework asks how the resulting behavioural dynamics combine over time to generate emergent dispersal. The two approaches are therefore complementary: mechanistic studies resolve the physical origin of movement, while data-driven generative models quantify its effective consequences at larger spatial and temporal scales.

The present model assumes that local swimming statistics remain approximately stationary and that individual trajectory fragments provide representative samples of the underlying locomotor process. These assumptions are reasonable during dilute exploratory swimming in the absence of imposed chemical gradients or fluid flow, but they will necessarily break down as environmental information becomes spatially structured. During host location, zoospores respond to chemical, electrical and physical cues through context-dependent changes in reorientation, retention and encystment (Kasteel *et al*., 2023, 2024). Recent studies have begun to identify receptors and signalling components involved in these responses (Pei *et al*., 2024; Ji *et al*., 2025), but much less is known about how receptor activation is translated into time-resolved changes in speed, turning and persistence. A natural extension of the present framework would therefore be to make behavioural transition rules explicitly dependent on local environmental variables such as chemical concentration, gradient direction or electrical potential.

The same principle applies to interactions among cells. Our simulations assume independent agents, an approximation that is appropriate at low zoospore densities but not necessarily in crowded populations. *Phytophthora* zoospores can form auto-aggregates through chemotaxis and bioconvection (Savory *et al*., 2014), while more general work on microswimmers shows that hydrodynamic interactions can alter individual trajectories and generate coordinated dynamics (Mo *et al*., 2025). Extending the model to include local cell density, secreted signals or fluid-mediated interactions would therefore provide a route from individual dispersal towards explicitly collective behaviour.

The obstacle experiments already illustrate how environmental structure can be incorporated without fundamentally altering the architecture of the model. The present environment is homogeneous except for impenetrable obstacles, but the same framework could accommodate increasingly heterogeneous representations of the zoospore habitat. Spatial fields could encode chemical or electrical gradients, local water flow, or host-derived signals, while reconstructed geometries could represent soil pores, air-water interfaces or root surfaces (Bassani *et al*., 2020; Yu *et al*., 2021). In such models, the behavioural rules governing the agent could remain explicit while becoming conditional on the local state of the environment. This would permit direct testing of how environmental complexity reshapes emergent dispersal without abandoning the data-driven structure of the model.

A further extension would be to move beyond motile unicellular agents towards growing filamentous organisms. Conceptually, many of the same modelling principles would apply to a growing hyphal tip: local rules would govern extension speed, directional persistence, arrest, reorientation and branching, while the trajectory of the advancing tip would leave behind a persistent structure rather than a transient path. The resulting architecture would therefore emerge from repeated local decisions in much the same way that dispersal emerges here from repeated swimming decisions. The experimental data required to infer such rules would of course differ substantially from those used for freely swimming zoospores and would require dedicated imaging and analysis strategies (Riquelme, 2013; Harris, 2025). Nevertheless, the same general framework (quantitative observation, local-rule inference, generative simulation and predictive challenge) could provide a basis for modelling filamentous growth as well as microbial dispersal.

In conclusion, we present a data-driven framework linking quantitative microscopy to agent-based modelling through the direct inference of local behavioural rules from experimental trajectories. The convergence between an empirical model and an independently inferred hidden Markov model supports the robustness of the behavioural organisation identified from the data, while predictive transfer to a different species and physical environment shows that these rules remain transferable across contexts. More broadly, our results illustrate how behavioural inference can bridge statistical description and generative modelling while preserving interpretability. The framework is not intended to reconstruct every molecular or biomechanical event underlying local motion, but rather to capture sufficient effective rules to reproduce and predict system-level behaviour. Its modular structure provides a natural basis for progressively incorporating environmental heterogeneity, collective interactions, host-associated signals and, ultimately, other forms of biological growth and dispersal. Predictive modelling of complex biological systems may therefore depend not only on increasing model sophistication, but on identifying local rules that are both experimentally grounded and sufficiently robust to support prediction under new conditions.

## Acknowledgments

We are indebted to Jean-Jacques Técourt, formerly a lecturer at the Classes Préparatoires aux Grandes Écoles (Lycée Masséna, France), for introducing the last author to the Caml Light programming language, and to François Graner (Université Paris Cité, France) for insightful discussions and valuable advice during the development of this project. We are grateful to Véronique Fritière and Linux Azur for the opportunity to present and discuss cellular automata implemented using an early version of the ABCA framework at the Journées Méditerranéennes du Logiciel Libre in 2011 (Sophia Antipolis, France). We thank Eric Galiana (Institut Sophia Agrobiotech, France) for providing the polystyrene microsphere suspension, and Tijs Ketelaar and Francine Govers (Wageningen University & Research, the Netherlands) for insightful discussions.

## Data availability

The time-lapse image stacks used for trajectory extraction and predictive validation in obstacle-filled environments, together with the corresponding TrackMate XML files, simulation outputs, and representative animated simulations, are available on Zenodo (doi.org/10.5281/zenodo.21898773). The source code of the ABCA framework, simulation plugins, Python analysis scripts, and calibrated model parameters used in this study are publicly available on GitHub (<u>github.com/EEvangelisti/ABCA</u>).

## Use of generative AI

Generative artificial intelligence was used to assist with software development and language editing. All AI-generated code, text, and suggestions were systematically reviewed, verified, and, where appropriate, modified by the authors before inclusion in the final software or manuscript. The authors take full responsibility for the accuracy and integrity of the work.

## Author contributions

**Joëlle Le Berre:** Investigation, Formal analysis, Data curation, Visualization, Writing - review & editing. **Agnès Attard:** Conceptualization, Investigation, Formal analysis, Supervision, Funding acquisition, Writing - review & editing.

**Edouard Evangelisti:** Conceptualization, Methodology, Software, Formal analysis, Visualization, Supervision, Project administration, Funding acquisition, Writing - original draft, Writing - review & editing.

## Notes

https://doi.org/10.5281/zenodo.21898773

### Competing Interest Statement

The authors have declared no competing interest.

### Summary of Updates

The manuscript has been substantially revised to broaden its conceptual framing, particularly in the Introduction and Discussion, and to emphasise the distinction between reproducing observed behaviour and predicting emergent dynamics beyond the conditions used to build the model. The results and conclusions remain unchanged.

### Funding

- Agence Nationale de la Recherche: ANR-11-LABX-0028-01, ANR-15-IDEX-01, ANR22-CE20-0021

