## Supporting Figures 1-10 for "Data-driven inference of local behavioural rules predicts emergent properties of *Phytophthora* zoospore dispersal"

### Contents

**Figure S1.** *Phytophthora nicotianae* trajectory length distribution. 2

**Figure S2.** Visual representation of individual *P. nicotianae* zoospore trajectories. 3

**Figure S3.** Relationship between swimming speed and acceleration. 4

**Figure S4.** Direct threshold-based state assignment produces fragmented behavioural episodes. 5

**Figure S5.** Hysteresis stabilises behavioural episodes. 6

**Figure S6.** Validation of the empirical agent-based model against experimental trajectories. 7

**Figure S7.** Visual representation of ABCA-simulated zoospore trajectories. 8

**Figure S8.** Selection of hidden Markov models describing zoospore swimming behaviour. 9

**Figure S9.** Biological interpretation of the two-state hidden Markov model. 10

**Figure S10.** Validation of the two-state hidden Markov model against experimental trajectories. 11

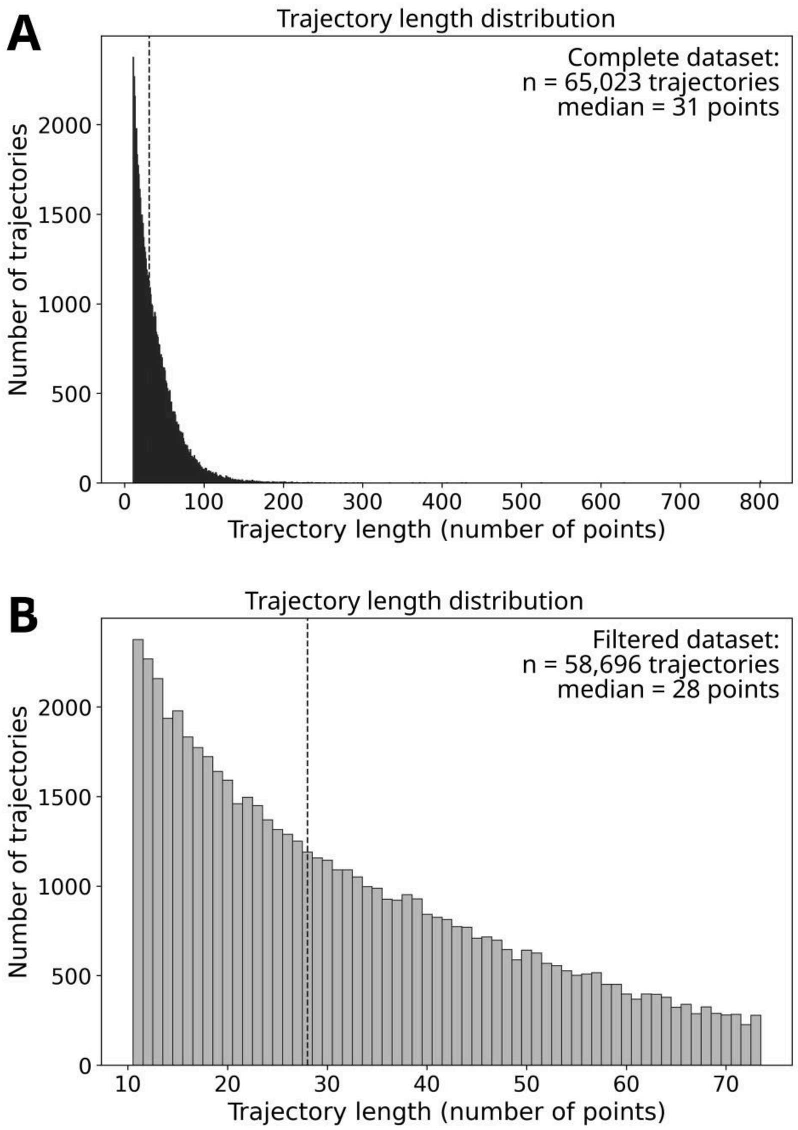


**Figure S1.** ***Phytophthora nicotianae*** **trajectory length distribution. (A-B)** A total of 65,023 *P. nicotianae* zoospore trajectories were obtained using TrackMate. Histograms show the complete dataset **(A)** and the dataset after excluding trajectories with lengths above the 90th percentile **(B)**. The latter was used for all subsequent quantitative analyses to reduce the influence of abnormally long trajectories, which are more likely to represent tracking artefacts or non-biological objects.

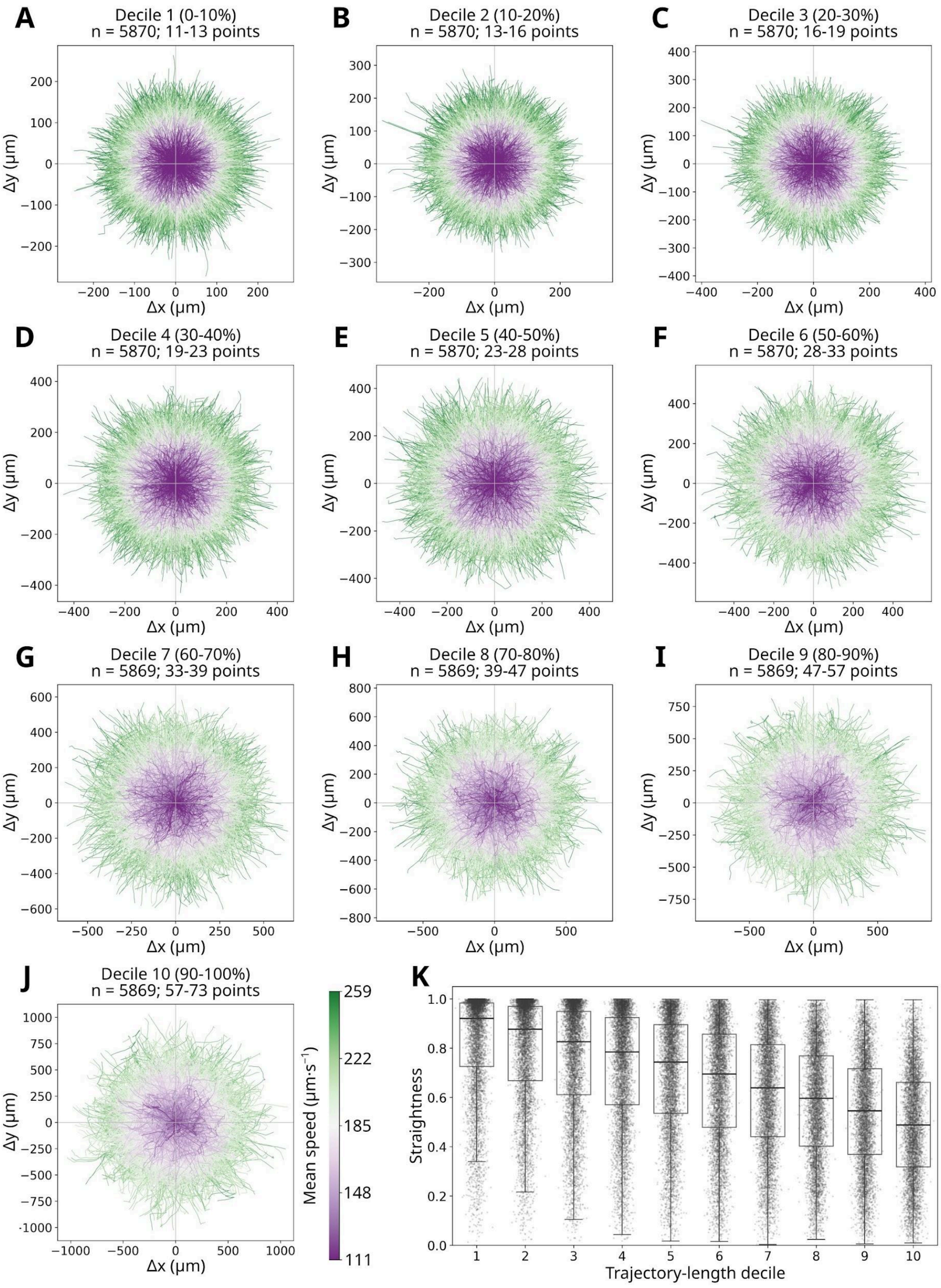


**Figure S2. Visual representation of individual *P. nicotianae* zoospore trajectories. (A-J)** Individual zoospore trajectories were grouped into deciles by length and translated so that all trajectories originated at (0, 0). Trajectories are coloured according to their mean swimming speed (µm/s), from low (purple) to high (green). Each decile is displayed using its own spatial scale to facilitate visual comparison of trajectory geometry within each length class. **(K)** Straightness of individual zoospore trajectories for each trajectory-length decile. Boxplots indicate the interquartile range, with the horizontal line representing the median. Dots represent individual straightness values.

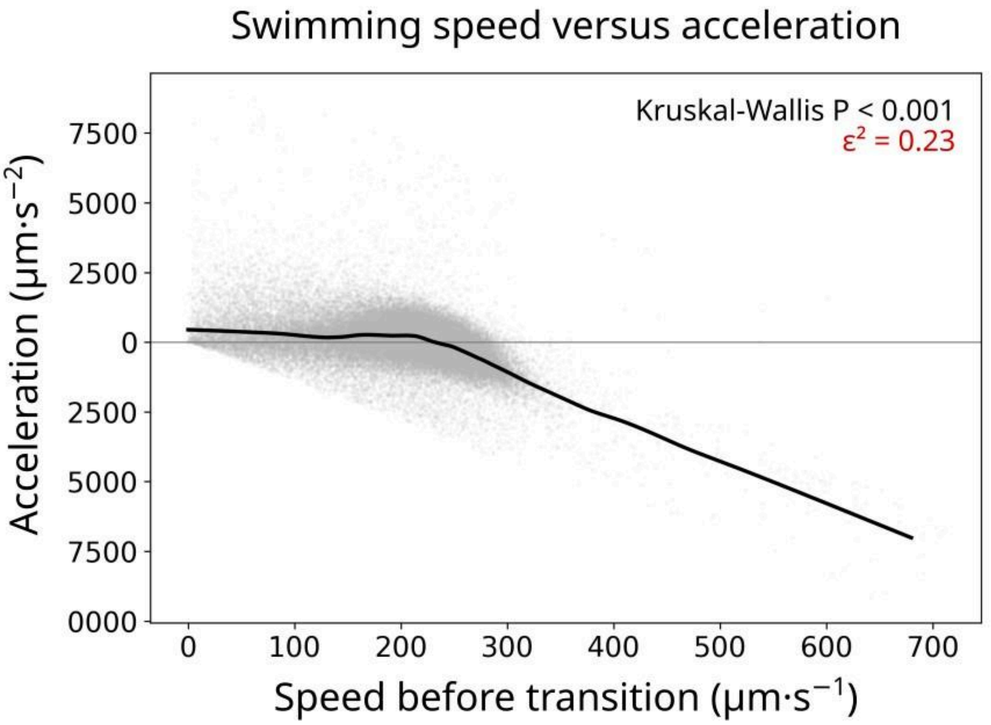


**Figure S3. Relationship between swimming speed and acceleration.** Relationship between swimming speed immediately before a transition and the corresponding instantaneous acceleration. Grey points represent individual observations, and the black curve shows a locally weighted regression (LOESS). Differences in acceleration distributions across speed deciles were assessed using a Kruskal-Wallis test ($P < 0.001$, $\varepsilon^2 = 0.23$).

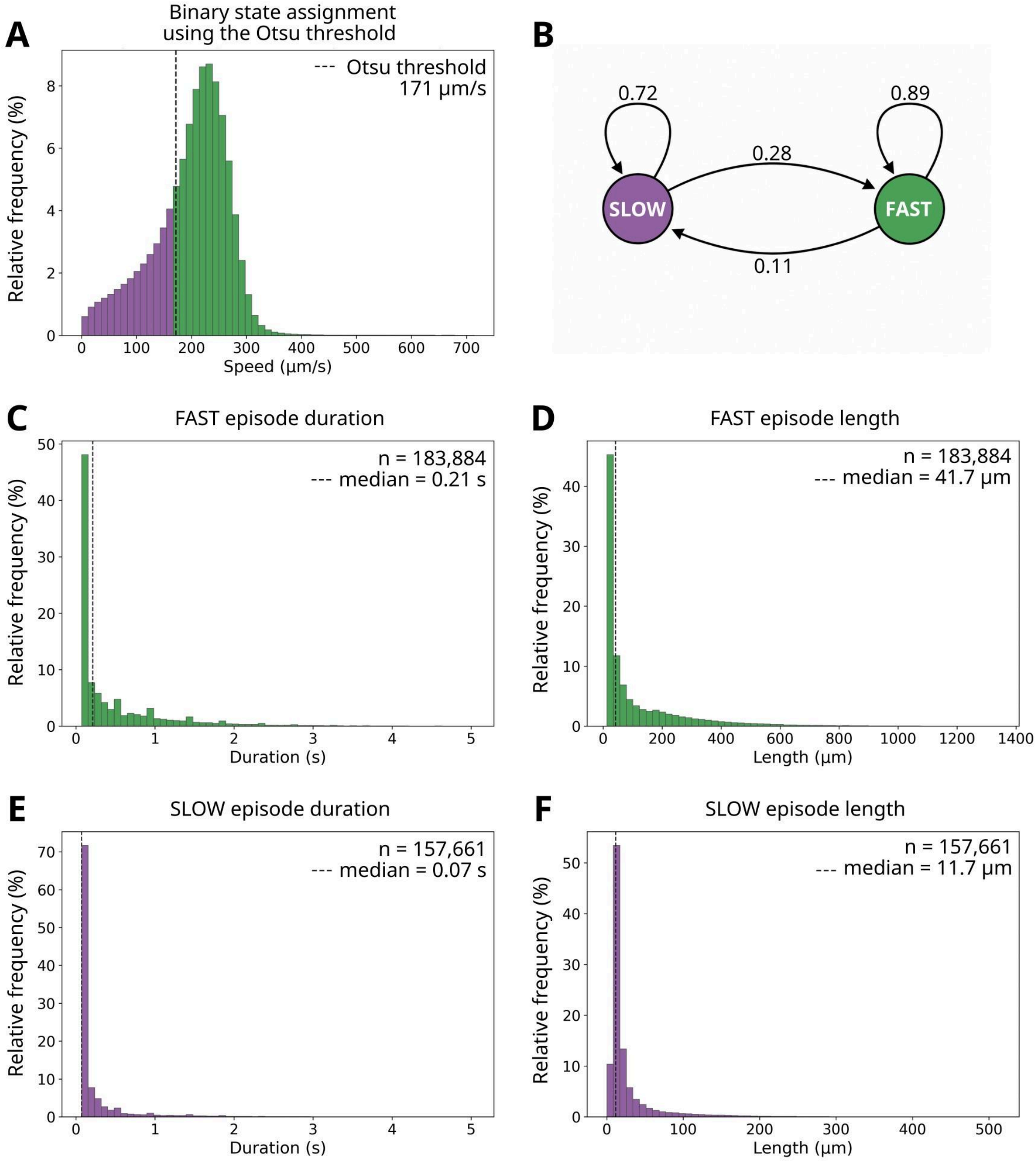


**Figure S4. Direct binary state assignment using the Otsu threshold produces fragmented behavioural episodes.** Swimming speeds were partitioned into SLOW and FAST states using the Otsu threshold without hysteresis. **(A)** Distribution of instantaneous swimming speeds after binary state assignment. The dashed line indicates the Otsu threshold separating SLOW (purple) and FAST (green) observations. **(B)** Transition probabilities estimated from the resulting binary state sequence. **(C, E)** Distributions of FAST- and SLOW-state episode durations. **(D, F)** Distributions of the corresponding episode path lengths. Direct frame-wise thresholding generated numerous very short behavioural episodes, resulting in frequent state transitions.

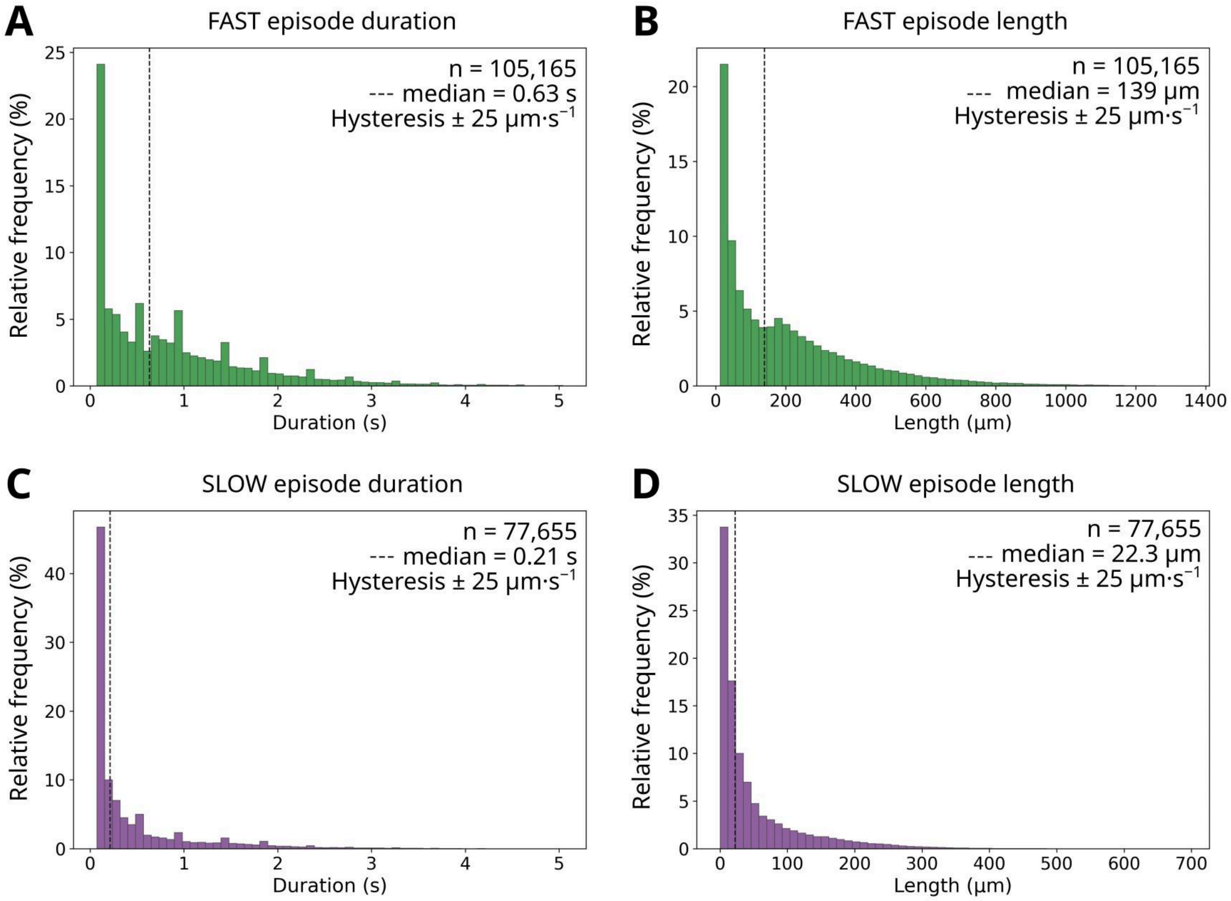


**Figure S5. Hysteresis stabilises behavioural episodes.** Swimming speeds were partitioned into SLOW and FAST states using the Otsu threshold with a hysteresis of ± 25 µm·s$^{-1}$. **(A, C)** Distributions of FAST and SLOW episode durations. **(B, D)** Distributions of the corresponding episode path lengths.

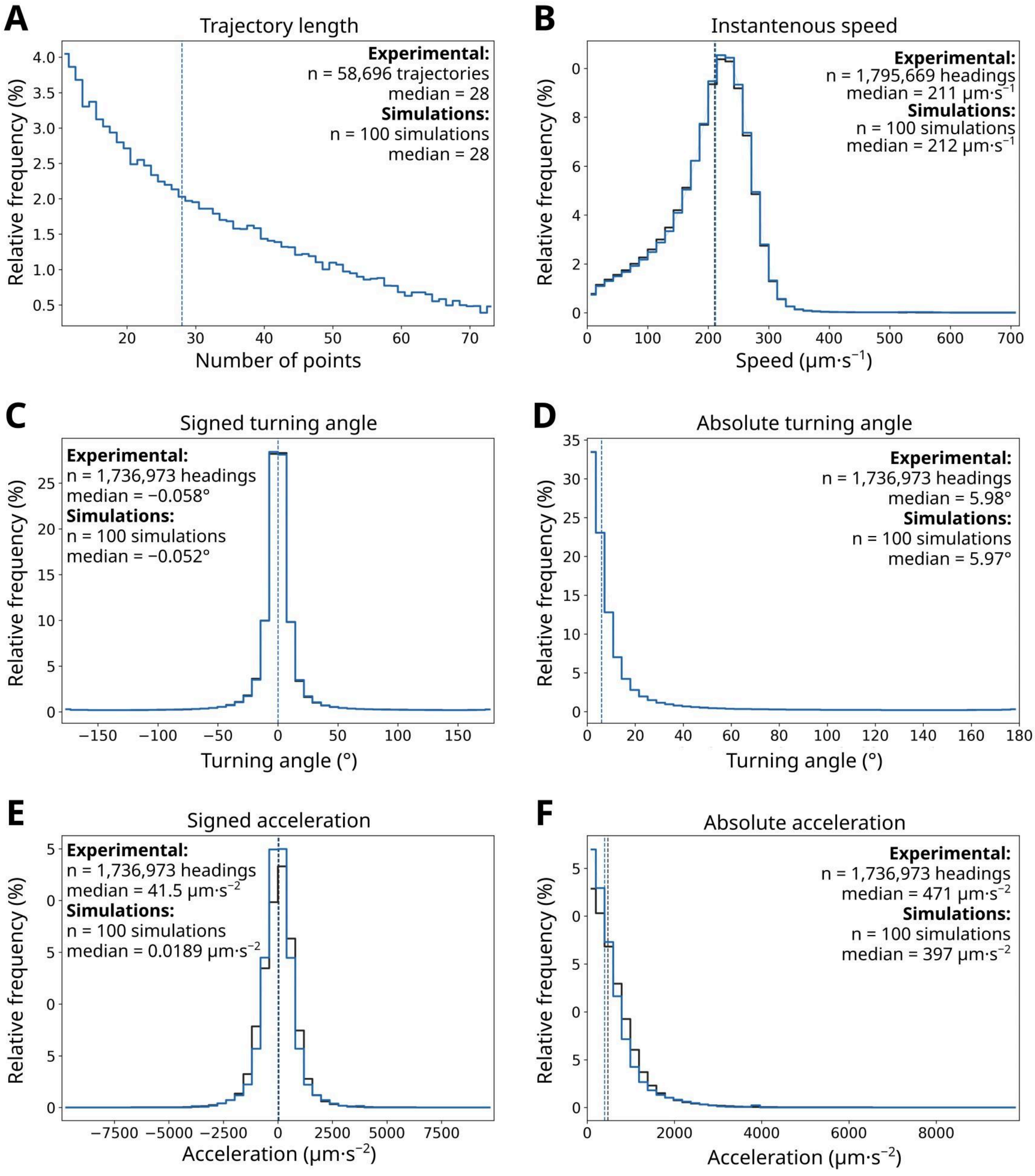


**Figure S6. Validation of the empirical agent-based model against experimental trajectories.** Experimental trajectories (black) were compared with an ensemble of 100 independent stochastic simulations generated with the empirical ABCA model (blue). **(A)** Simulated trajectories were resampled to match the experimental trajectory-length distribution before computing trajectory descriptors. **(B)** Instantaneous swimming speed. **(C)** Signed turning angle. **(D)** Absolute turning angle. **(E)** Signed acceleration. **(F)** Absolute acceleration. Vertical dashed lines indicate the medians of the experimental (black) and simulated (blue) distributions.

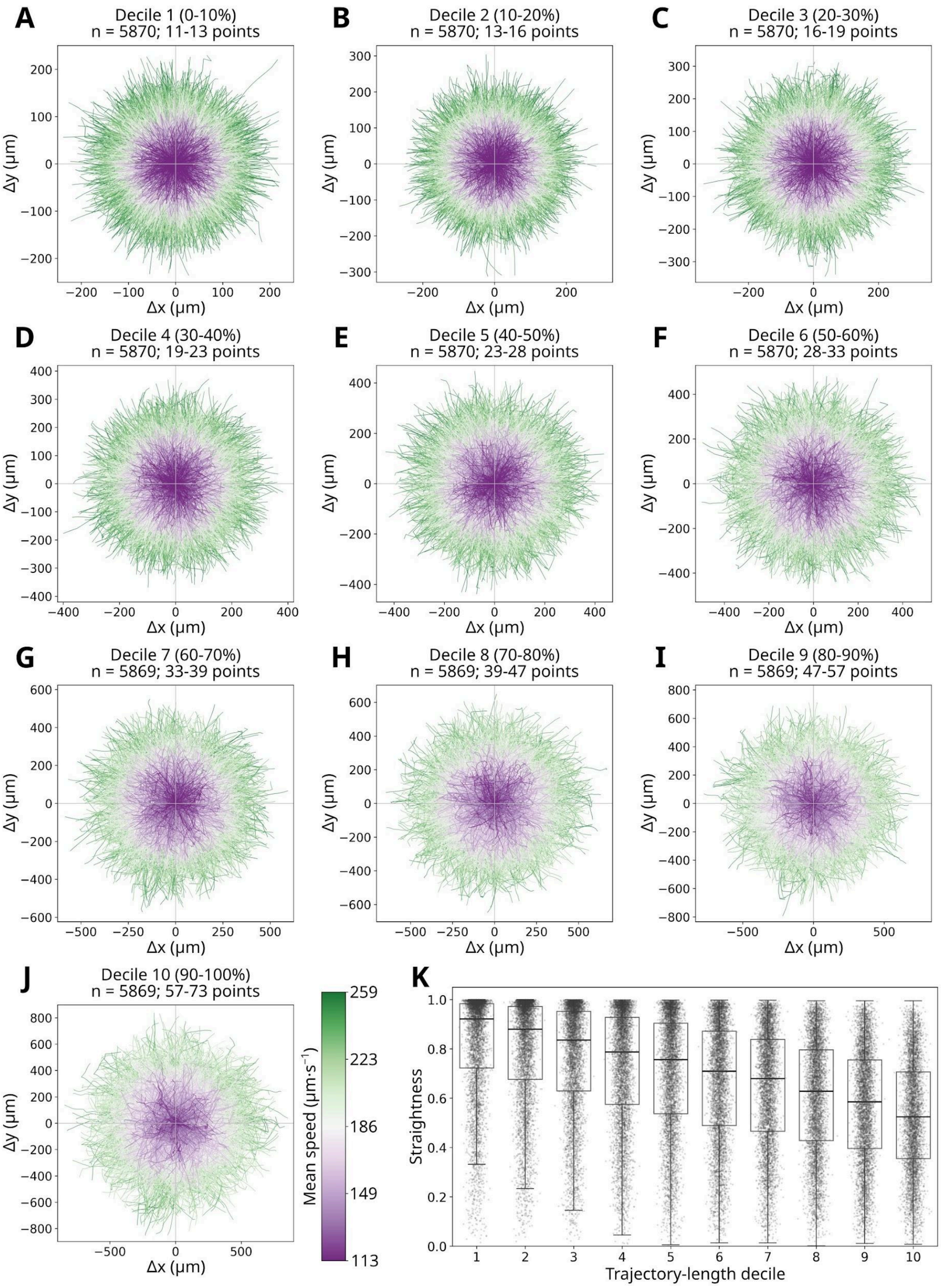


**Figure S7. Visual representation of ABCA-simulated zoospore trajectories.** This figure was generated using the same analysis pipeline and graphical representation as Figure S2. **(A-J)** Individual simulated zoospore trajectories were grouped into deciles by length and translated so that all trajectories originated at (0, 0). Trajectories are coloured according to their mean swimming speed (μm/s), from low (purple) to high (green). Each decile is displayed using its own spatial scale to facilitate visual comparison of trajectory geometry within each length class. **(K)** Straightness of individual zoospore trajectories for each trajectory-length decile. Boxplots indicate the interquartile range, with the horizontal line representing the median. Dots represent individual straightness values.

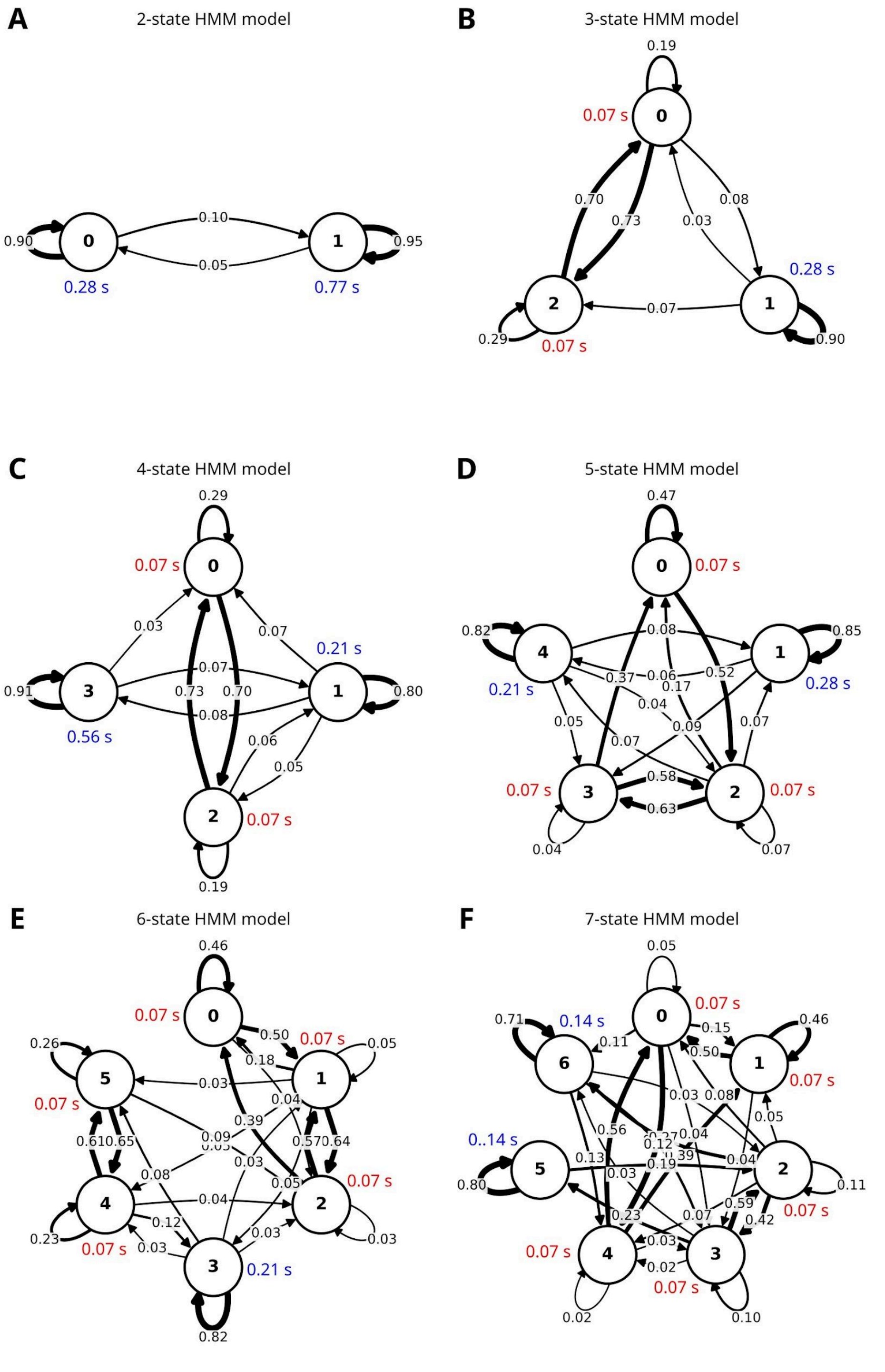


**Figure S8. Selection of hidden Markov models describing zoospore swimming behaviour. (A-F)** Transition graphs of the best-fitting HMMs containing two **(A)**, three **(B)**, four **(C)**, five **(D)**, six **(E)**, and seven **(F)** hidden states. Circles represent hidden states, and directed arrows represent state transitions. Numbers indicate transition probabilities. Coloured numbers indicate the median sojourn time (s) in each state; red values correspond to the minimum observable duration of a single experimental frame (0.07 s). Arrow thickness is proportional to transition probability. Self-loops represent state persistence.

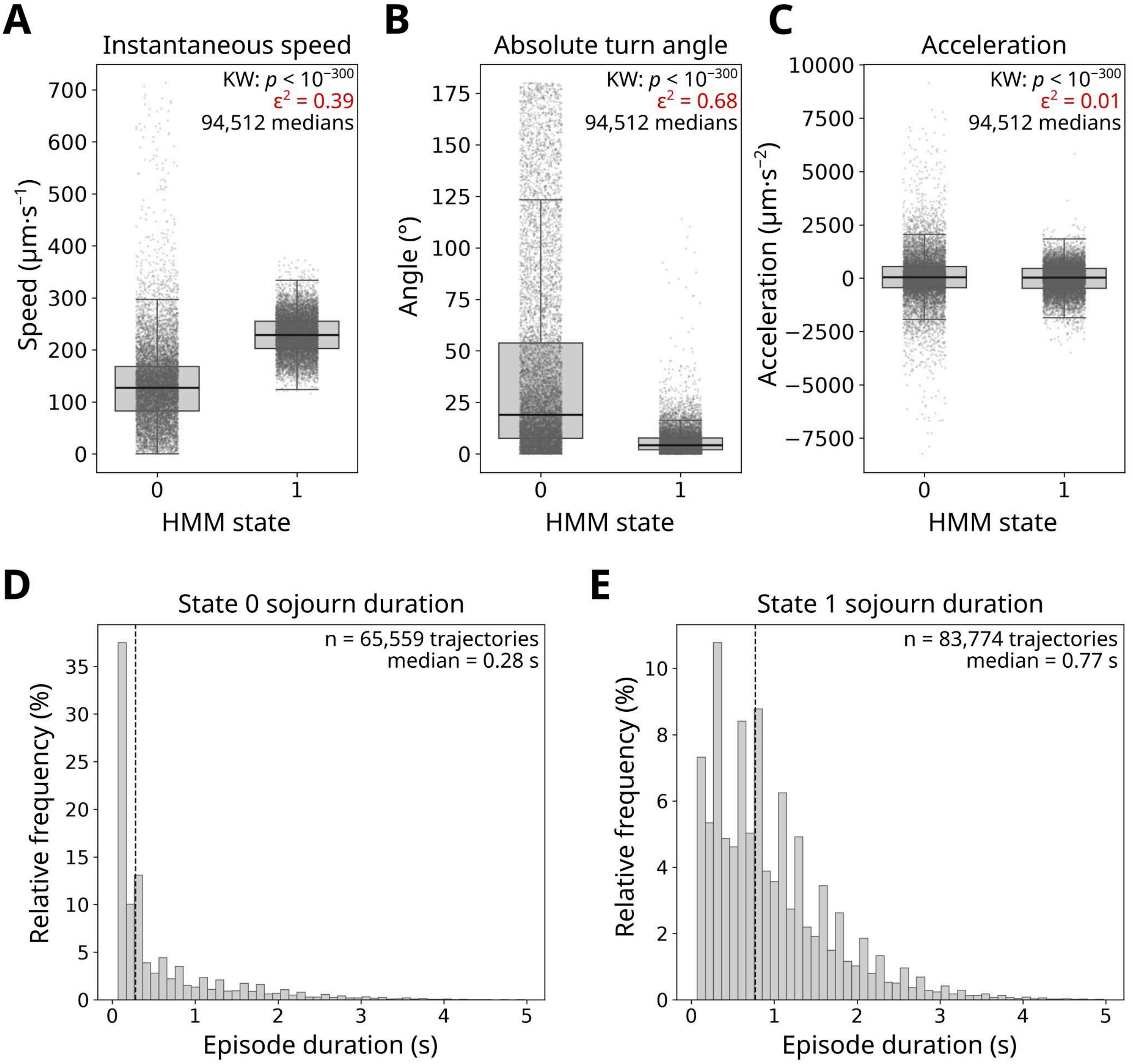


**Figure S9. Biological interpretation of the two-state hidden Markov model. (A-C)** Biological characteristics of the two decoded HMM states. For each trajectory and decoded state, a single median value was computed to avoid treating successive observations as independent replicates. Boxplots show the distribution of trajectory medians, while grey dots represent individual trajectory medians (10,000 randomly selected medians displayed to prevent clutter). Statistical differences between states were assessed using Kruskal-Wallis (KW) tests, and effect sizes were reported as epsilon squared ($\varepsilon^2$, red). The two latent states differ strongly in instantaneous swimming speed (**A**; $\varepsilon^2 = 0.39$) and absolute turning angle (**B**; $\varepsilon^2 = 0.68$), whereas acceleration differs only weakly between states despite remaining statistically significant (**C**; $\varepsilon^2 = 0.01$). **(D-E)** Distributions of episode durations (state sojourn times) for the two decoded states. Histograms show the relative frequency of continuous occupancy episodes, and dashed vertical lines indicate median episode durations. State 0 exhibits short occupancy episodes (median = 0.28 s), whereas state 1 exhibits substantially longer episodes (median = 0.77 s).

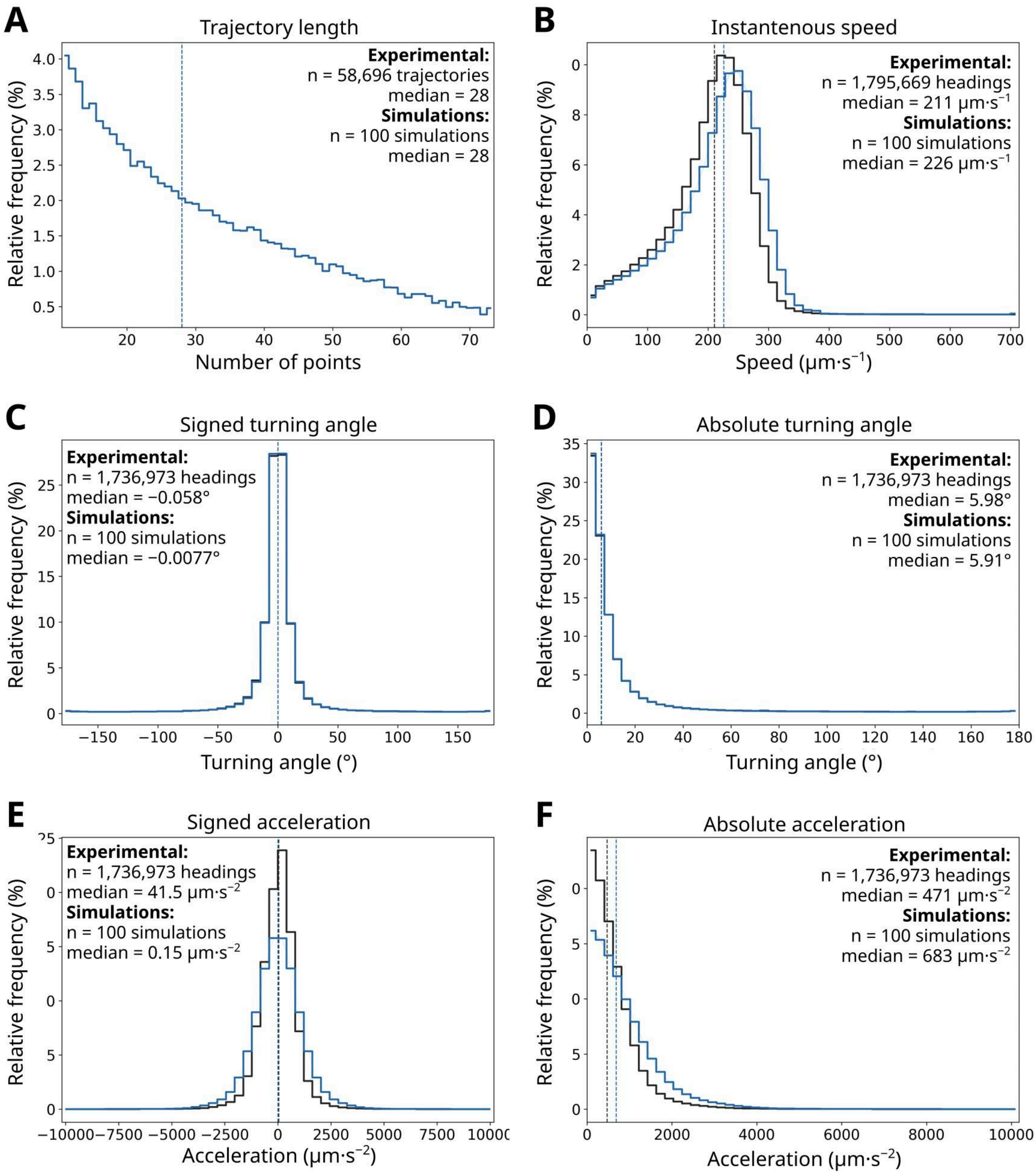


**Figure S10. Validation of the two-state hidden Markov model against experimental trajectories.** Experimental trajectories (black) were compared with an ensemble of 100 independent stochastic simulations generated using the two-state hidden Markov model (blue), providing an independent assessment of the behavioural model inferred from trajectory data. **(A)** Simulated trajectories were resampled to match the experimental trajectory-length distribution before computing local trajectory descriptors. **(B)** Instantaneous swimming speed. **(C)** Signed turning angle. **(D)** Absolute turning angle. **(E)** Signed acceleration. **(F)** Absolute acceleration. Vertical dashed lines indicate the medians of the experimental (black) and simulated (blue) distributions.
